# IGF2 mediates Hippo signaling to control liver size

**DOI:** 10.1101/2024.10.27.620455

**Authors:** Zhenxing Zhong, Ruxin Jin, Yiting Zhong, Li Zhang, Deqian Chen, Zhihan Jiao, Fanhui Zhou, Rui Zhu, Jian Wu, Rui Dong, Kuiran Dong, Fei Lan, Yu Wang, Kun-Liang Guan, Fa-Xing Yu

**Author notes:** Corresponding author: (FL), (YW), (FXY). Equal contribution (Zhenxing Zhong, Ruxin Jin, Yiting Zhong).

## Abstract

The Hippo pathway is a central mechanism in organ size control, but the mediator for its function remains elusive. Here, we show that the expression of insulin-like growth factor II (*IGF2*) is directly induced by YAP/TAZ transcription cofactors of the Hippo pathway in a developmental stage- and cell type-specific manner. In mouse livers, *Igf2* expression is sustained by YAP/TAZ in hepatoblasts and immature hepatocytes at fetal and neonatal stages, coupling with rapid cell proliferation and liver size expansion, whereas turned off in matured hepatocytes following YAP/TAZ inactivation to prevent liver overgrowth. In contrast, YAP/TAZ fails to regulate *Igf2* expression in cholangiocytes and diverse liver mesenchymal cells where epigenetic barriers, including DNA and histone methylation, are implanted near *Igf2* promoters. Furthermore, IGF2 activates IGF1R signaling and promotes normal and neoplastic liver growth, while the inactivation of IGF2 or IGF1R effectively blocks YAP/TAZ-induced hepatomegaly and hepatoblastoma. In conclusion, our findings reveal that IGF2 is a *bona fide* mediator of the Hippo pathway, indicating a fundamental role of the Hippo-IGF2-IGF1R signaling axis in organ size regulation and malignancies.

## INTRODUCTION

The precise regulation of organ size is fundamental to development and homeostasis, but how organs know when to stop growing has been an enigma for centuries ^1,2^. The Hippo pathway has recently been identified as a central regulator of organ size in multicellular organisms ^3–6^. In mammals, this pathway is typically represented as a kinase cascade in which the activity of Yes-associated protein (YAP) and transcriptional co-activator with PDZ-binding motif (TAZ, also known as WWTR1), two major downstream effectors, is restricted by upstream kinases and adaptor proteins ^5–12^. In mice, ectopic expression of YAP or inactivation of its upstream regulators leads to dramatic enlargement of affected organs ^5–10^. Moreover, YAP and TAZ are considered proto-oncoproteins, with their aberrant activation frequently observed in cancer specimens ^13–18^. Therefore, the Hippo pathway is a critical mechanism regulating both normal and neoplastic growth, participating in organ size control and tumorigenesis.

As transcription co-factors, YAP and TAZ are speculated to control tissue growth by modulating the expression of genes involved in cell proliferation, differentiation, and survival. However, until now, no known target gene of YAP/TAZ has been demonstrated to regulate organ size under physiological conditions ^19–22^. It also remains unclear how YAP/TAZ modulates gene expression to drive oncogenic transformation and progression. Hence, identifying key mediators of YAP/TAZ, if any, is crucial for understanding how the Hippo pathway regulates normal and neoplastic growth. Based on previous studies, we reason that such a downstream mediator should fulfill several criteria: (i) it is directly induced by YAP/TAZ; (ii) it is likely a growth factor; (iii) its expression is dynamically associated with organ growth; and (iv) it can be reactivated during regeneration and tumorigenesis.

Among known growth factors, insulin-like growth factor II (IGF2) has garnered our attention. In developing mouse livers, the expression of *Igf2* is dynamic and tightly correlated with cell proliferation and YAP/TAZ activity, as revealed by previous single-cell RNA sequencing (scRNA-seq) data (Fig S1A) ^23^. Under physiological conditions, *Igf2* is predominantly expressed in fetal, neonatal, and placental tissues, where it promotes developmental organ growth ^24–26^. *Igf2* expression sharply declines around two weeks after birth, becoming nearly undetectable in adult mice ^27^. However, the molecular mechanisms responsible for the postnatal silencing of *Igf2* remain unclear. In humans, excessive *IGF2* expression causes Beckwith-Wiedemann syndrome, characterized by pronounced body overgrowth, whereas its deficiency results in Russell-Silver syndrome, marked by severe growth retardation ^28^. Consistently, mice with heterozygous *Igf2* deletion exhibit approximately a 60% reduction in body weight, while *Igf2* overexpression significantly enlarges embryos and multiple organs ^25,29–31^. Moreover, elevated *IGF2* expression is observed in various cancer specimens and is closely associated with poor prognosis ^32–36^. These findings underscore the importance of IGF2 in organ size regulation and cancer, promoting us to explore its potential crosstalk with the Hippo pathway.

In this study, we show that YAP/TAZ directly regulates *IGF2* expression in a developmental stage- and cell type-specific manner. By activating insulin-like growth factor 1 receptor (IGF1R) signaling, IGF2 acts as a *bona fide* mediator of the Hippo pathway and is indispensable for regulating organ size and tumorigenesis. Moreover, the IGF2-IGF1R signaling axis can be targeted to modulate organ size and to treat cancers driven by YAP/TAZ activation.

## RESULTS

### The Hippo pathway directly regulates *IGF2* expression

The mouse liver is an ideal model for studying organ size regulation. Immediately after birth, liver parenchymal cells proliferate rapidly to expand the liver size. By 2-3 weeks old, the growth rate gradually decelerates to a basal level, and a relatively constant liver-to-body weight ratio (∼4%) is maintained throughout life ^4,10,37^. We had established mice with liver-specific deletion (*Albumin-Cre*, *Alb-Cre*) of upstream regulators in the hippo pathway and analyzed their effects on liver size during postnatal development ^9,10^. Strikingly, *Sav1^−/−^;Wwc1*^−/−^*;Wwc2*^−/−^ (referred to as *Sav1^−/−^;Wwc*^−/−^) mice exhibited rapid hepatomegaly without evident tumor growth ^9,10^. To identify critical downstream alterations in these livers, we performed transcriptome analysis on livers of 3-week-old *Sav1^−/−^;Wwc*^−/−^ mice, which displayed more than 4-fold increases in liver-to-body weight ratios (Figs 1A, B). YAP/TAZ was activated in *Sav1^−/−^;Wwc*^−/−^ livers, as indicated by the increased expression of canonical YAP/TAZ target genes, including cysteine-rich protein 61 (*Cyr61*) and connective tissue growth factor (*Ctgf*), which encode extracellular matrix proteins known to regulate cell adhesion and migration (Figs 1C, D and S1B, C) ^19–21^.

**Figure 1.**
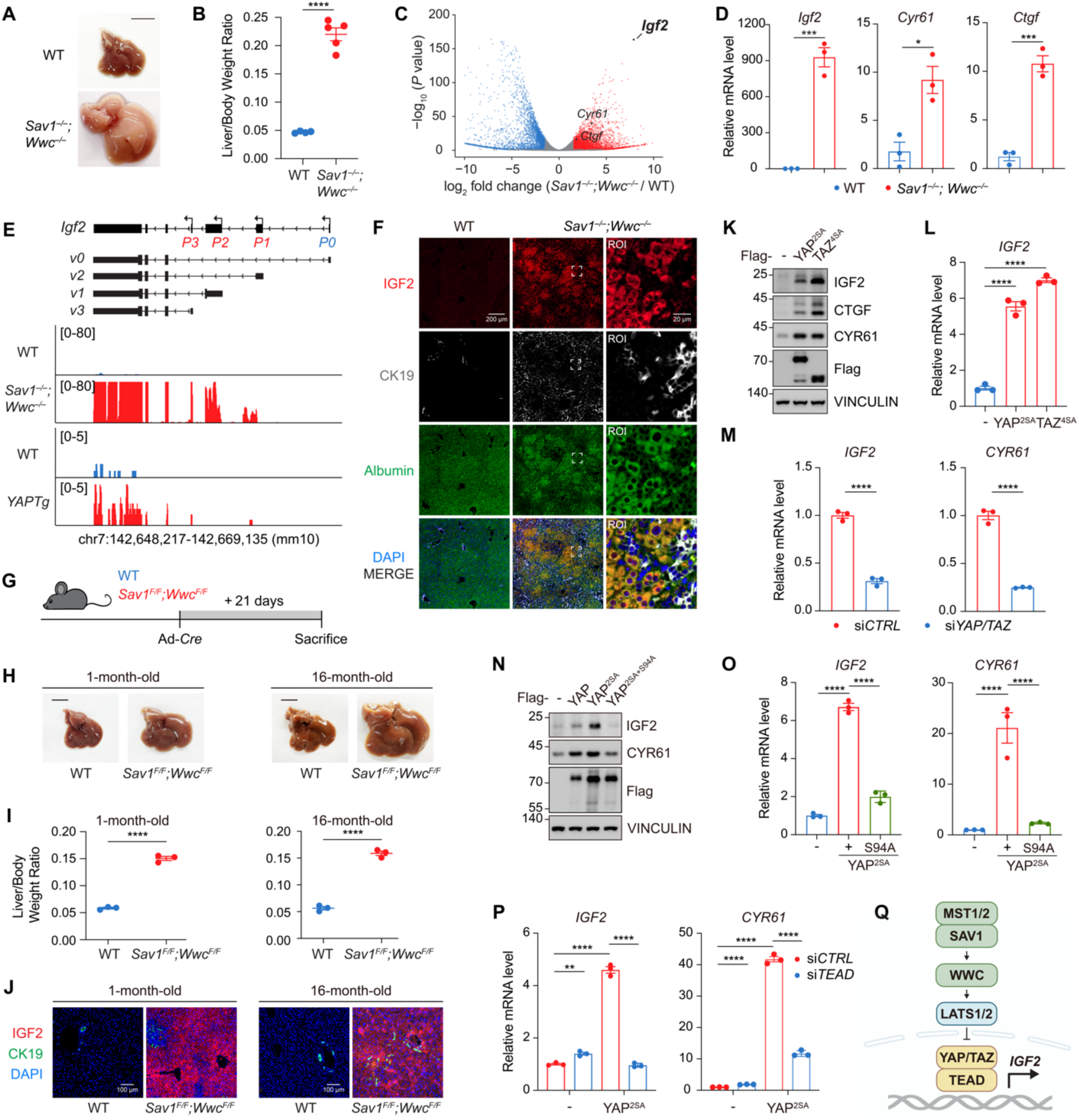
The expression of *IGF2* is directly regulated by the Hippo pathway. (A and B) Liver-specific deletion of *Sav1* and *Wwc1/2* in mice results in rapid hepatomegaly. Gross liver images (A) and liver-to-body weight ratios (B) of 3-week-old wild-type (WT) and *Sav1^−/−^;Wwc1*^−/−^*;Wwc2*^−/−^ (referred to as *Sav1^−/−^;Wwc*^−/−^ for simplicity) mice are shown. Scale bar in A: 1 cm. (C) Volcano plot exhibiting differentially expressed genes in 3-week-old *Sav1^−/−^;Wwc*^−/−^ livers compared to WT livers, with genes of interest marked. (D) Quantitative RT-PCR results of *Igf2*, *Cyr61*, and *Ctgf* mRNA levels in 3-week-old WT and *Sav1^−/−^;Wwc*^−/−^ livers. (E) Genomic tracks displaying RNA-seq reads at the *Igf2* gene locus, comparing *Sav1^−/−^;Wwc*^−/−^ livers (upper) and *YAPTg* livers (lower) with age-matched WT livers. The RNA-seq data for *ApoE-rtTA-YAP* transgenic (*YAPTg*) livers were analyzed using data from GSE178227. (F) Immunostaining of IGF2 (red), CK19 (white), Albumin (green), and nuclei (blue) in 3-week-old WT and *Sav1^−/−^;Wwc*^−/−^ livers. Scale bar: 200 μm. Region of interest (ROI) scale bar: 20 μm. (G-J) Acute deletion in *Sav1^F/F^;Wwc1^F/F^;Wwc2^F/F^*(*Sav1^F/F^;Wwc^F/F^*) mice via Ad-*Cre* injection at different ages (1 or 16 months old) leads to rapid liver enlargement and robust IGF2 upregulation. Schematic diagram (G), gross liver images (H), liver-to-body weight ratios (I), and immunostaining (J) of IGF2 (red), CK19 (green), and nuclei (blue) are shown. Scale bars: 1 cm (H), 100 μm (J). (K and L) Overexpression of YAP^2SA^ and TAZ^4SA^ (constitutively active forms) in HEPG2 cells induces *IGF2* expression. Immunoblotting (K) and quantitative RT-PCR (L) results are shown. (M) Knockdown of *YAP/TAZ* in HEPG2 cells impairs *IGF2* expression. (N and O) The S94A mutation in YAP^2SA^, which disrupts TEAD binding, attenuates its ability to induce *IGF2* and *CYR61* expression. Immunoblotting (N) and quantitative RT-PCR (O) results are shown. (P) Knockdown of *TEAD1/3/4* in HEPG2 cells impairs the activity of YAP^2SA^ in upregulating *IGF2* and *CYR61* expression. (Q) Schematic diagram illustrating that the Hippo pathway regulates *IGF2* expression in a YAP/TAZ- and TEAD-dependent manner. Data in B, D, I, L, M, O, and P are presented as mean ± SEM from at least three independent biological replicates. Each point in B, D, and I represents an individual mouse. *P* values in B, D, I, L, M, O, and P were assessed by unpaired Student’s t-tests. *p < 0.05, **p < 0.01, ***p < 0.001, ****p < 0.0001.

Notably, we observed a nearly 1000-fold increase of *Igf2* mRNA levels in 3-week-old *Sav1^−/−^;Wwc*^−/−^ livers (Figs 1C, D). The *Igf2* gene is transcribed into multiple isoforms by different promoters (Fig 1E) ^24–27^. The exon-specific reads revealed that the fetal *P1*, *P2*, and *P3* promoters were activated in *Sav1^−/−^;Wwc*^−/−^ livers, while the placenta-specific *P0* promoter remained quiescent (Fig 1E). The elevated *IGF2* expression in *Sav1^−/−^;Wwc*^−/−^ livers was also confirmed by immunostaining (Fig 1F). Similar results were observed in the liver of 1-month-old *ApoE-rtTA-YAP* transgenic (*YAPTg*) mice, another frequently used YAP activation model (Figs 1E and S1B, C) ^6,22^. To avoid potential confounding factors during liver development, we induced acute gene deletion by intravenous injection of *Cre*-expressing adenovirus (Ad-*Cre*) into *Sav1^F/F^;Wwc1^F/F^;Wwc2^F/F^* mice (Fig 1G). Intriguingly, *Sav1* and *Wwc1/2* ablation in mice at different ages (1 or 16 months old) led to rapid liver enlargement with a profound upregulation of *IGF2* expression (Figs 1H-J and S1B-D). Conversely, genetic inactivation of *Yap/Taz* notably suppressed *Igf2* expression in 1-week-old mouse livers, a developmental stage in which *Igf2* was highly expressed (Figs S1B-C).

The expression of IGF2 appeared to be restricted to hepatocytes with Albumin expression (Fig 1F). To further investigate whether IGF2 alterations in these livers were regulated by the Hippo pathway in a cell-autonomous manner, we isolated primary hepatocytes from mice and induced YAP/TAZ activation *in vitro* (Fig S1E). We found that *Igf2* was upregulated upon *Sav1* and *Wwc1/2* deletion, but this induction was significantly reduced when *Yap/Taz* was knocked down (Figs S1F, G). Moreover, in human hepatoblastoma HEPG2 cells, overexpression of YAP^2SA^ and TAZ^4SA^ (constitutively active forms) or deletion of upstream kinases *LATS1/2* also induced *IGF2* expression (Figs 1K, L and S1H-J). On the contrary, YAP/TAZ deficiency or LATS1/2 overexpression significantly impaired the expression of *IGF2* in HEPG2 cells (Figs 1M and S1H, J). These results indicate that the Hippo pathway directly regulates *IGF2* expression.

YAP/TAZ interacts with DNA-binding transcription factors, particularly TEA domain transcription factors (TEAD1-4, collectively referred to as TEAD), to regulate gene transcription ^38–40^. In HEPG2 cells, we found that a S94A mutation of YAP (TEAD-binding defective) significantly decreased its ability to induce *IGF2* expression (Figs 1N, O). Similar results were observed in cells with *TEAD1/3/4* knockdown (Fig 1P). Moreover, the knockdown of *Tead1-4* in primary mouse hepatocytes with *Sav1* and *Wwc1/2* deficiency also impaired the expression of *Igf2* (Fig S1K). Together, these results suggest that *IGF2* expression is regulated by the Hippo pathway in a YAP/TAZ- and TEAD-dependent manner (Fig 1Q).

### YAP/TAZ induces *IGF2* expression in immature hepatocytes and hepatoblastomas

To further assess the effect of the Hippo pathway on *Igf2* expression and liver growth, we analyzed histological and molecular changes in 3-week-old mouse livers with deletions of one or more Hippo pathway genes (*Nf2*, *Sav1*, *Wwc1/2*, or compound knockout) (Fig 2A) ^9,10^. The expression of *Igf2* remained largely unchanged in *Nf2^−/−^*, *Sav1^−/−^*, or *Wwc^−/−^* livers with only mild increases in YAP/TAZ activity and liver size, but was robustly induced in *Sav1^−/−^;Wwc*^−/−^ livers with moderate YAP/TAZ activation and elevated liver-to-body weight ratios (Fig 2A). Unexpectedly, *Igf2* expression was only slightly induced in *Nf2^−/−^;Wwc1^+/–^;Wwc2^+/–^*(referred to as *Nf2^−/−^;Wwc*^+/–^) livers, despite these livers exhibiting the highest YAP/TAZ activity and liver-to-body weight ratios (Fig 2A). Embryonic activation of YAP/TAZ in the liver triggers polarized differentiation towards cholangiocytes ^10,41,42^. Indeed, *Nf2^−/−^;Wwc*^+/–^ livers were predominantly occupied by cholangiocytes (CK19+), but these cholangiocytes did not express *Igf2* (Fig 2A). In contrast, *Sav1^−/−^;Wwc*^−/−^ livers retained a large number of hepatocytes (Albumin+) and showed increased expression of *Afp* and *Gpc3*, genes expressed in hepatoblasts or liver progenitors in fetal and neonatal livers (Fig 2A) ^43–45^. These results indicate that *Igf2* expression is associated with not only YAP/TAZ activity but also lineage specifications during liver development.

**Figure 2.**
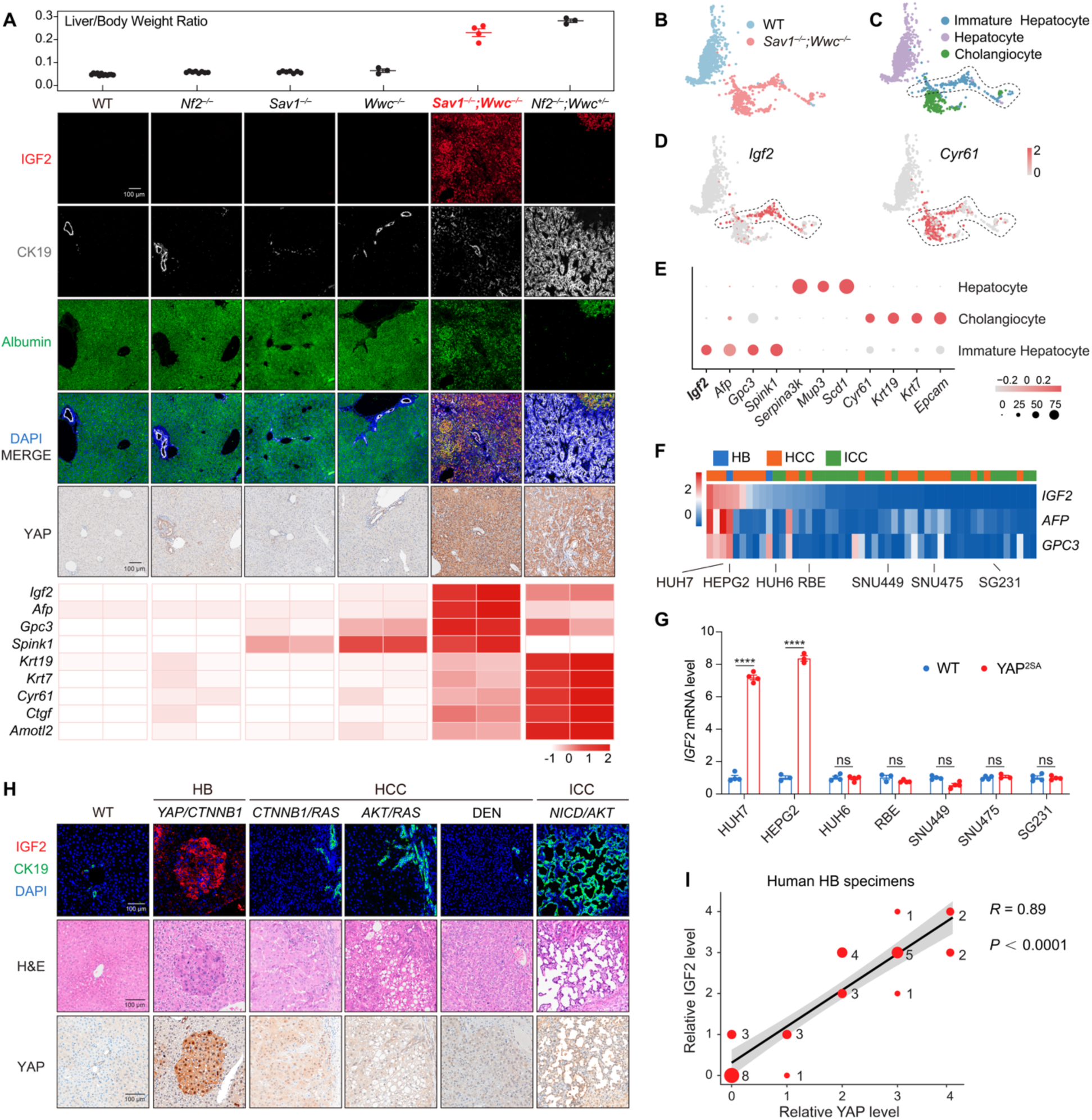
IGF2 is induced by YAP/TAZ specifically in immature hepatocytes and hepatoblastomas. (A) *Igf2* expression is associated with YAP/TAZ activity and lineage specification during liver development. Upper panel: Liver-to-body weight ratios of 3-week-old mouse livers with *Nf2*, *Sav1*, *Wwc1/2*, or compound knockout. Middle panel: Immunostaining images of IGF2 (red), CK19 (white), Albumin (green), nuclei (blue), and YAP (IHC). Lower panel: Expression profiles of *Igf2*, immature hepatocyte markers (*Afp*, *Gpc3*, *Spink1*), cholangiocyte markers (*Krt19*, *Krt7*), and canonical YAP/TAZ target genes (*Cyr61*, *Ctgf*, *Amotl2*). The transcriptome was analyzed using RNA-seq data from mouse livers with *Nf2*, *Sav1*, *Wwc1/2*, or compound deletions (CRA008364). Scale bar: 100 μm. (B and C) UMAP representation of integrated liver parenchymal cells, generated by combining scRNA-seq data from 3-week-old WT (GSE171993) and *Sav1^−/−^;Wwc*^−/−^ livers. Cells are colored by samples (B) or cell types (C). (D) Expression profiles of *Igf2* and *Cyr61* in liver parenchymal cells. (E) Expression of selected markers for liver parenchymal cells. Dot size and color indicate the proportion of cells expressing marker genes in each cluster and the average gene expression level, respectively. (F) Transcriptome analysis from the CCLE dataset indicates that *IGF2* is highly expressed in liver cancer cells with characteristics of immature hepatocytes. HB: hepatoblastoma; HCC: hepatocellular carcinoma; ICC: intrahepatic cholangiocarcinoma. (G) YAP^2SA^ effectively induces *IGF2* expression in HEPG2 and HUH7 cells but not in cells where *IGF2* is silenced. Quantitative RT-PCR results are presented as mean ± SEM from at least three independent biological replicates. *P* values were assessed by unpaired Student’s t-tests. ****p < 0.0001; ns: not significant. (H) IGF2 is specifically expressed in YAP/TAZ-activated HB driven by *YAP/CTNNB1* co-expression. H&E staining and immunostaining of IGF2 (red), CK19 (green), nuclei (blue), and YAP (IHC) are shown. Scale bar: 100 μm. (I) *IGF2* expression closely correlates with YAP/TAZ activity in human HB specimens. Quantitative analysis of immunostaining results for IGF2 and YAP (Fig S2N). Dot size indicates the number of specimens. Correlation coefficients were calculated using Spearman’s correlation.

To further characterize the spatiotemporal regulation of *Igf2* expression by the Hippo pathway, we performed scRNA-seq analysis on 3-week-old WT and *Sav1^−/−^;Wwc*^−/−^ livers (Figs 2B and S2A). Unsupervised clustering of 10,316 cells by uniform manifold approximation and projection (UMAP) identified 11 cell types according to the expression of canonical markers (Figs S2B-D). Three clusters of liver parenchymal cells were defined: hepatocytes, cholangiocytes, and a group of immature hepatocytes that appeared in *Sav1^−/−^;Wwc*^−/−^ livers (Figs 2C-E). These immature hepatocytes resembled neonatal hepatocytes (P1-P7) during postnatal development, as evidenced by high expression of *Afp* and *Gpc3* (Figs S2E-G). Notably, *Igf2* was highly expressed in these immature hepatocytes (Figs 2D, E and S2C, D). However, *Igf2* was absent in cholangiocytes, fibroblasts, and endothelial cells, even though YAP/TAZ was active in these cells, as indicated by high *Cyr61* expression (Figs 2D, E and S2C, D). Moreover, *Igf2* was also undetectable in matured hepatocytes and other non-parenchymal cells where YAP/TAZ was inactive (Figs 2D, E and S2C, D). These results suggest that YAP/TAZ specifically regulates *Igf2* expression in immature hepatocytes.

Next, we investigated whether the cell type-specific regulation of *IGF2* by the Hippo pathway is conserved in liver cancers. YAP/TAZ activation has been observed in three subtypes of liver cancer: hepatoblastoma (HB), hepatocellular carcinoma (HCC), and intrahepatic cholangiocarcinoma (ICC) ^13,14,16–18^. By analyzing the Cancer Cell Line Encyclopedia (CCLE) dataset, we found that *IGF2* was expressed in some HB cells and a small subset of HCC cells, but undetectable in all ICC cells (Fig 2F). In most cancer cells with *IGF2* expression, *AFP* and *GPC3* were also highly expressed (Fig 2F). Interestingly, ectopic expression of YAP effectively induced *IGF2* in liver cancer cells with basal *IGF2* expression, including HEPG2 and HUH7 cells (Figs 2F, G and S2H-J). In contrast, in cells where *IGF2* was silenced, active YAP failed to promote *IGF2* expression, although canonical YAP/TAZ targets, such as *CYR61* and *CTGF*, were robustly induced (Figs 2F, G and S2H-J). Hence, YAP/TAZ may regulate *IGF2* expression only in liver cancer cells exhibiting characteristics of immature hepatocytes.

Activation of YAP/TAZ in mouse livers induces tumorigenesis ^8–10,12^. In a collection of mouse liver tumors induced by inactivating different Hippo pathway genes, we noticed that IGF2 was highly expressed in HCC but not in ICC or bile duct hamartomas surrounding HCC (Figs S2K, L). Transcriptome analysis revealed that these HCC induced by dysregulated Hippo pathway exhibited immature features, as evidenced by high *Afp* and *Gpc3* expression (Fig S2M). In mice, hydrodynamic injection (HDI) of oncogenes into hepatocytes or treatment with carcinogen diethylnitrosamine (DEN) causes different liver tumors. Interestingly, *IGF2* expression was robust in the HB model (*YAP*/*CTNNB1*) but not in other tumor models such as HCC (*CTNNB1*/*RAS*, *AKT*/*RAS*, or DEN) or ICC (*NICD*/*AKT*), even though YAP was slightly induced in these tumors (Fig 2H) ^13,46,47^. These observations prompted us to investigate the role of YAP on *IGF2* expression in HB, which originates from immature hepatocytes during embryonic and neonatal stages and frequently exhibits dysregulation of the Hippo pathway ^13,14^. We collected tissues from 33 human HB patients and quantitatively assessed the expression of YAP and IGF2 in tumors by immunostaining (Fig S2N). Notably, a strong correlation (R=0.89) was observed between YAP activation and *IGF2* expression in these HB specimens (Fig 2I). Collectively, these findings suggest that IGF2 is highly expressed in immature hepatocytes and HB, which exhibit neonatal hepatoblast-like characteristics and moderate YAP/TAZ activation.

### Promoter DNA methylation prevents YAP/TAZ-induced *IGF2* expression

*IGF2* expression in fetal and neonatal livers is driven by several fetal promoters (Fig 3A). Unlike canonical YAP/TAZ target genes, *IGF2* is induced by YAP/TAZ only in specific cell types (Fig 2). We then set to identify the mechanisms underlying this cryptic regulation. DNA methylation typically results in gene silencing and is a pivotal epigenetic mechanism regulating dynamic gene expression during development ^48^. In various liver cancer cells, *IGF2* expression is inversely correlated with the levels of DNA 5-methylcytosine (5mC) at its fetal promoters, as indicated by Reduced-Representation Bisulfite Sequencing (RRBS) data from the CCLE dataset and targeted methylation sequencing (Figs 3B and S3A) ^49^. In liver cancer cells with hypermethylated *IGF2* fetal promoters, treatment with decitabine, a potent DNA methyltransferase inhibitor (DNMTi), reduced 5mC levels and reactivated *IGF2* transcription (Figs 3C and S3B-C). Interestingly, in the presence of DNMTi, the transcription of *IGF2* was further enhanced by YAP overexpression (Figs 3C and S3B-C). These results indicate that promoter DNA methylation is a barrier to preventing YAP/TAZ-induced *IGF2* transcription.

**Figure 3.**
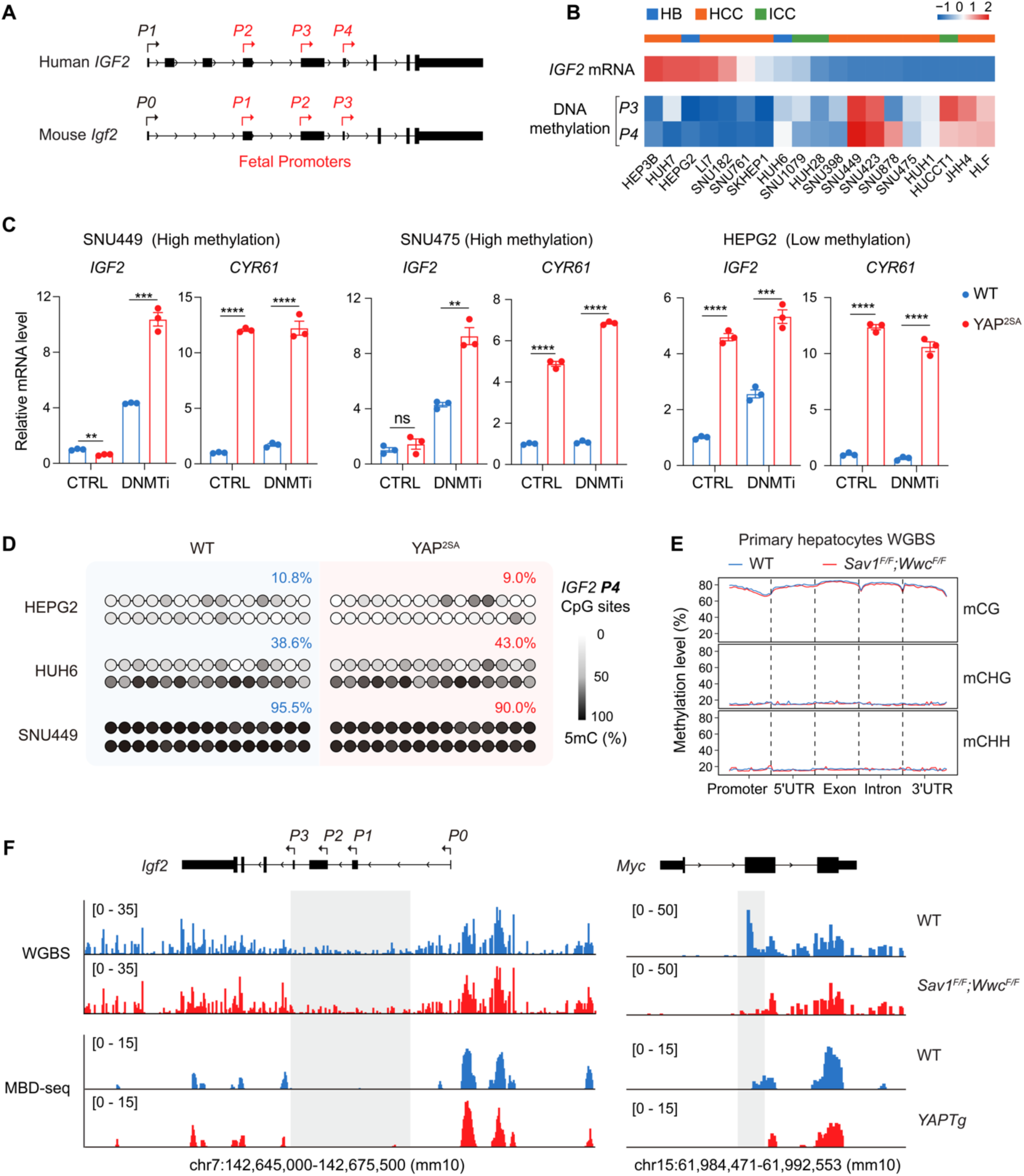
Promoter DNA methylation is a barrier to preventing YAP/TAZ-induced *IGF2* transcription. (A) Schematic diagram illustrating the promoters of human *IGF2* and mouse *Igf2* genes, with fetal promoters marked in red. (B) Expression and reduced representation bisulfite sequencing (RRBS) data from the CCLE dataset indicate that *IGF2* expression is inversely correlated with 5-methylcytosine (5mC) levels at its *P3/P4* fetal promoters. HB: hepatoblastoma; HCC: hepatocellular carcinoma; ICC: intrahepatic cholangiocarcinoma. (C) In SNU449 and SNU475 cells with hypermethylated promoters, *IGF2* expression is reactivated and further enhanced by YAP^2SA^ when treated with the DNA methyltransferase inhibitor (DNMTi) decitabine. Data are presented as mean ± SEM from at least three independent biological replicates. *P* values were assessed by unpaired Student’s t-tests. **p < 0.01, ***p < 0.001, ****p < 0.0001; ns: not significant. (D) YAP^2SA^ expression has no obvious effect on DNA methylation at the *IGF2 P4* promoter. Each dot represents a CpG site within the *IGF2 P4* region, and the color reflects the 5mC level at that site. (E) Whole-genome bisulfite sequencing (WGBS) results indicate no overt difference between WT and *Sav1^F/F^;Wwc*^F/F^ hepatocytes on global DNA methylation across different genomic regions. Blue lines: WT hepatocytes; red lines: *Sav1^F/F^;Wwc^F/F^* hepatocytes. (F) Genomic tracks displaying WGBS and methylated DNA immunoprecipitation sequencing (MBD-seq, GSE178227) reads. The *Igf2* fetal promoters (*P1/P2/P3*) are hypomethylated while the placental promoter (*P0*) is hypermethylated. Deletion of *Sav1;Wwc1/2* (*Sav1^F/F^;Wwc^F/F^*) or YAP transgenic expression (*YAPTg*) has minimal effect on DNA methylation of *Igf2* promoters (left). In contrast, the *Myc* locus, shown as a positive control, exhibits partial demethylation upon YAP activation, consistent with previous reports (right).

Recently, YAP has been reported to induce an oncogenic transcriptional program through Ten-Eleven Translocation methylcytosine dioxygenase 1 (TET1)-mediated epigenetic remodeling ^22^. We then asked whether YAP activation leads to active remodeling of DNA methylation near *IGF2* promoters. However, the effect of active YAP on DNA demethylation of *IGF2* promoters was minimal in multiple liver cancer cells (Figs 3D and S3D, E). To further assess the *in vivo* effects of YAP activation on DNA methylation, we used AAV8-*TBG-Cre* to induce hepatocyte-specific YAP activation in *Sav1^F/F^;Wwc^F/F^* mice (Fig S3F). Primary hepatocytes were isolated 21 days post-injection and subjected to RNA-seq and whole-genome bisulfite sequencing (WGBS) (Fig S3G). Transcriptome profiling confirmed YAP activation, elevated *Igf2* expression, and an immature state of these primary hepatocytes with *Sav1* and *Wwc1/2* knockout (Fig S3H). Consistent with a previous report, while YAP activation led to regional DNA demethylation in certain genes such as *Myc*, global DNA methylation changes were minimal (Figs 3E, F and S3I, J) ^22^. For *Igf2* gene, the fetal promoters (*P1/2/3*) were hypomethylated, but there was no significant difference between WT and hepatocytes deficient in *Sav1* and *Wwc1/2* (Figs 3F and S3K). This pattern was also observed in *YAPTg* livers (Fig 3F) ^22^. These findings suggest that although regional 5mC modifications are barriers to YAP/TAZ-induced *IGF2* expression, activation of YAP/TAZ does not actively remove DNA methylation near *IGF2* promoters.

### Chromatin compaction impedes YAP/TAZ-induced *IGF2* expression

Although DNMTi treatment promoted *IGF2* transcription in several liver cancer cells, the extent of reactivation remained limited (Figs 3C and S3C). This suggests that additional epigenetic mechanisms may contribute to *IGF2* transcription and responsiveness to YAP activation. YAP activation alone failed to induce *IGF2* expression in SNU449 cells effectively; we treated this cell line with various histone-modifying enzyme inhibitors but observed no dramatic changes in *IGF2* transcription (Figs S4A-C). However, when combined with DNMTi, several inhibitors robustly boosted *IGF2* transcription (Figs S4A-C). Notably, EPZ6438 and GSK343, two inhibitors of Enhancer of Zeste Homolog 2 (EZH2), showed the most robust effect (Fig S4B). EZH2 is the catalytic component of Polycomb Repressive Complexes (PRC2), which primarily methylates histone H3 on lysine 27 (H3K27me3) to repress gene expression via chromatin compaction (Fig S4D) ^50^. Using chromatin immunoprecipitation (ChIP) analysis, we found that in cells with low *IGF2* expression, particularly ICC cells, *IGF2* promoters exhibited high levels of H3K27me3 (Figs 4A and S4E, F). The EZH2 inhibitor (EZH2i) led to H3K27me3 depletion at *IGF2* promotes and enhanced YAP-induced *IGF2* expression when combined with DNMTi in SNU449 cells (Figs 4B, C). In the presence of DNMTi and EZH2i, YAP effectively induced *IGF2* transcription in multiple cell lines with low or no basal expression, including HEK293A cells (Figs S4G-K). These results indicate that promoter histone methylation may be another barrier preventing YAP/TAZ-induced *IGF2* transcription.

**Figure 4.**
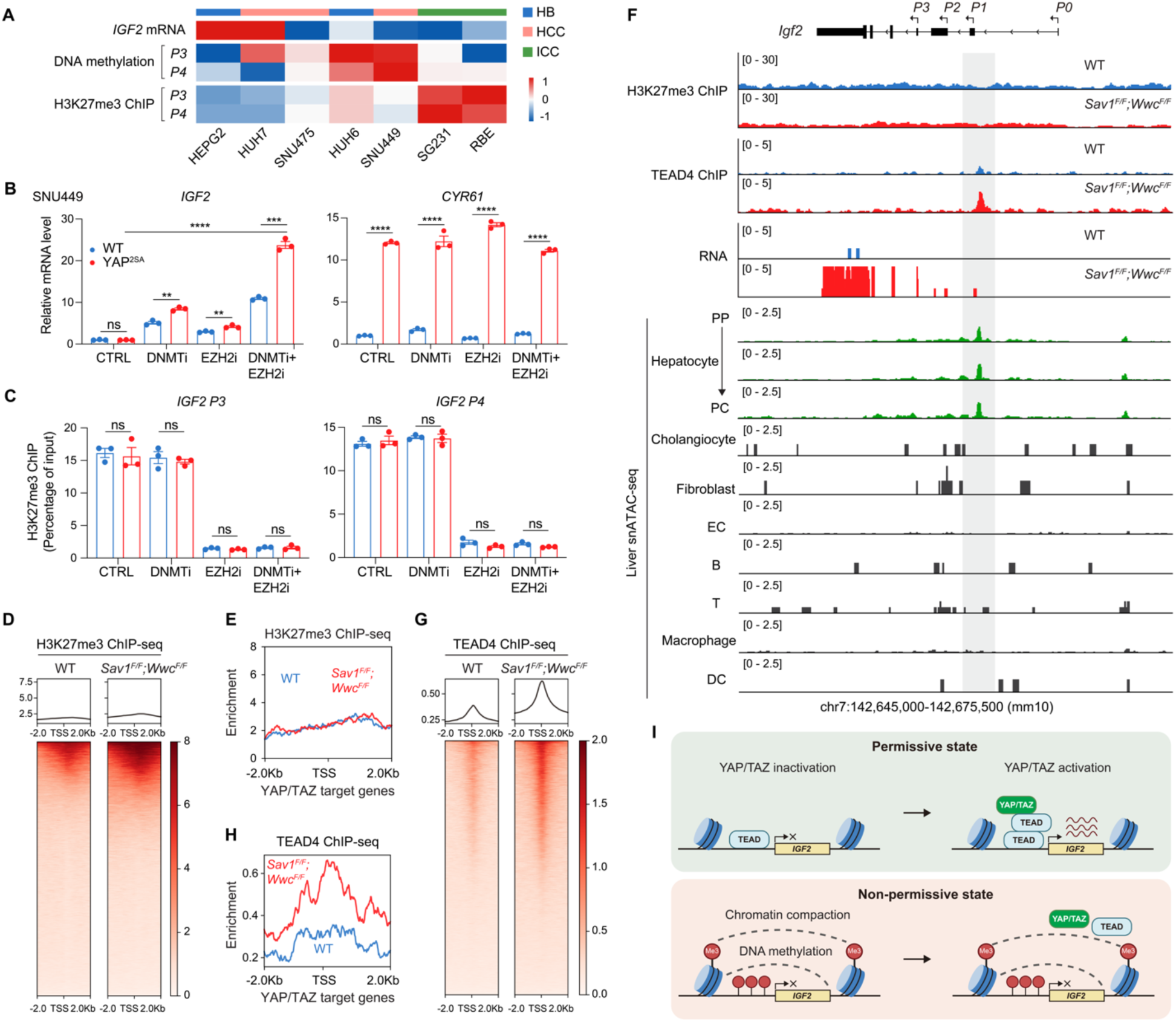
Chromatin compaction is another barrier impeding YAP/TAZ-induced *IGF2* transcription. (A) *IGF2* expression is inversely correlated with DNA 5mC and H3K27me3 levels near its *P3/4* fetal promoters. HB: hepatoblastoma; HCC: hepatocellular carcinoma; ICC: intrahepatic cholangiocarcinoma. (B) Treatment with the EZH2 inhibitor (EZH2i) EPZ6438, in combination with the DNA methyltransferase inhibitor (DNMTi) decitabine, enhances YAP-induced *IGF2* expression (left), while exerting minimal effect on *CYR61* expression (right). Data are presented as mean ± SEM from at least three independent biological replicates. *P* values were assessed by unpaired Student’s t-tests. **p < 0.01, ***p < 0.001, ****p < 0.0001; ns: not significant. (C) EPZ6438 treatment leads to H3K27me3 depletion near *IGF2 P3/P4* fetal promotes. Chromatin immunoprecipitation (ChIP) signals were normalized to input DNA and are presented as mean ± SEM. *P* values were assessed by unpaired Student’s t-tests; ns: not significant. (D) Heatmaps representing the enrichment of H3K27me3 ChIP-seq reads centered at transcription start sites (TSS). (E) Enrichment profiles representing the H3K27me3 ChIP-seq reads centered at TSS of canonical YAP/TAZ target genes in WT and *Sav1^F/F^;Wwc^F/F^* hepatocytes. Blue lines: WT hepatocytes; red lines: *Sav1^F/F^;Wwc^F/F^* hepatocytes. (F) Genomic tracks displaying H3K27me3 ChIP-seq, TEAD4 ChIP-seq, RNA-seq, and mouse liver single-nucleus ATAC-seq (snATAC-seq, González-Blas et al. 2024) reads at the *Igf2* locus. TEAD-binding sites are marked in gray, which are accessible only in hepatocytes. PP: periportal; PC: pericentral; EC: endothelial cell; DC: dendritic cell. (G) Heatmaps representing the enrichment of TEAD4 ChIP-seq (E) reads centered at TSS. (H) Enrichment profiles representing the TEAD4 ChIP-seq reads centered at TSS of canonical YAP/TAZ target genes in WT and *Sav1^F/F^;Wwc^F/F^* hepatocytes. Blue lines: WT hepatocytes; red lines: *Sav1^F/F^;Wwc^F/F^*hepatocytes. (I) Schematic diagram illustrating the permissive state (upper) and non-permissive state (lower) near *IGF2* promoters. In the permissive state, such as in hepatocytes, YAP/TAZ activation induces *IGF2* expression in a TEAD-dependent manner. In the non-permissive state caused by intense H3K27me3 or 5mC modifications, YAP/TAZ activation fails to induce *IGF2* expression due to these epigenetic barriers.

Next, we set to investigate the role of YAP on histone methylation. As indicated by ChIP analysis, while EZH2i robustly reduced H3K27me3 levels near *IGF2* promoters, the effect of active YAP was marginal (Figs 4C and S4E-F). We also performed ChIP sequencing (ChIP-seq) in primary mouse hepatocytes and observed no evident impact of YAP activation on global H3K27me3 levels (Figs 4D, E). In addition, H3K27me3 at *Igf2* promoters was low and minimally affected by YAP activation (Fig 4F). These results suggest that YAP/TAZ does not actively remove promoter H3K27me3 marks to induce *Igf2* expression. On the other hand, in YAP-activated hepatocytes, TEAD4 occupancy at *Igf2* fetal promoters was robustly increased, likely due to elevated TEAD4 expression upon YAP/TAZ activation (Figs 4F-H and S4L) ^22^. In hepatocytes, the basal levels of DNA and histone methylation at *Igf2* fetal promoters were low, allowing the binding of YAP/TAZ-TEAD complex and the induction of gene transcription (Figs 3F and 4F). This was further supported by single-nucleus ATAC-seq (snATAC-seq) data of adult mouse livers ^51^, in which the TEAD-binding site at *Igf2* fetal promoters was accessible in hepatocytes, but closed in cholangiocytes and different mesenchymal cells (Fig 4F). By contrast, the TEAD-binding sites at *Cyr61* and *Ctgf* promoters were accessible in multiple cell types (Fig S4M).

Together, these findings suggest that the responsiveness of *IGF2* expression to YAP/TAZ activation depends on the chromatin state of *IGF2* promoters (Fig 4I). In hepatocytes or cells in a permissive state, YAP/TAZ activation can effectively induce *IGF2* expression. Conversely, intense H3K27me3 or 5mC modifications create a non-permissive state near *IGF2* promoters in non-hepatocytes or some liver cancer cells, preventing YAP/TAZ-induced *IGF2* expression.

### IGF2 is required for YAP-induced hepatomegaly and tumorigenesis

As a growth factor critical for fetal and neonatal development, IGF2 is closely associated with organ growth ^25,28^. To investigate whether IGF2 is required for YAP/TAZ-induced liver growth and tumorigenesis, we generated an *Igf2* conditional knockout mouse line by introducing two LoxP sequences between exon 7 and exon 8, encompassing the functional coding sequence shared by all *Igf2* transcripts (Fig S5A). Liver-specific knockout mice (*Igf2^−/−^*, *Sav1^−/−^;Wwc*^−/−^, and *Igf2^−/−^;Sav1^−/−^;Wwc*^−/−^) were established using *Alb-Cre* (Figs 5A, B). Previous studies have shown that global *Igf2* deficiency in mice leads to a robust reduction in the size of multiple organs, including the liver ^25,52^. However, conditional knockout of *Igf2* in the liver alone had minimal effect on liver growth (Figs 5A-C). We reasoned that during early development, IGF2 produced by other organs might compensate for its loss in the liver via a systematic effect ^24,26,27^. In contrast, compared to the dramatically enlarged livers in *Sav1^−/−^;Wwc*^−/−^ mice, *Igf2* ablation (*Igf2^−/−^;Sav1^−/−^;Wwc*^−/−^) significantly reduced liver size, with the liver-to-body weight ratios decreased by approximately 70% (Figs 5A, C). As a mitogenic factor, IGF2 may induce liver growth by promoting cell proliferation. Indeed, expression of Ki67, a nuclear marker of dividing cells, was induced throughout the liver in *Sav1^−/−^;Wwc*^−/−^ livers but sharply decreased in *Igf2^−/−^;Sav1^−/−^;Wwc*^−/−^ livers (Figs 5A, D). Hepatocyte-specific 5-ethynyl-2’-deoxyuridine (EdU) incorporation assay showed consistent results (Figs 5E and S5B, C). To determine the molecular alterations after genetic inactivation of *Igf2*, we performed RNA-seq on these livers. As shown by principal component analysis (PCA) and clustering analysis, the global transcription profile was dramatically perturbed in *Sav1^−/−^;Wwc*^−/−^ livers but normalized in *Igf2^−/−^;Sav1^−/−^;Wwc*^−/−^ livers (Figs 5F and S5D). Intriguingly, in *Igf2^−/−^;Sav1^−/−^;Wwc*^−/−^ livers, the expression of immature hepatocyte markers, cholangiocyte markers, and proliferation markers was reduced to basal levels (Figs 5G and S5E). These results underscore the requirement of IGF2 for liver growth and hepatomegaly driven by YAP/TAZ activation.

**Figure 5.**
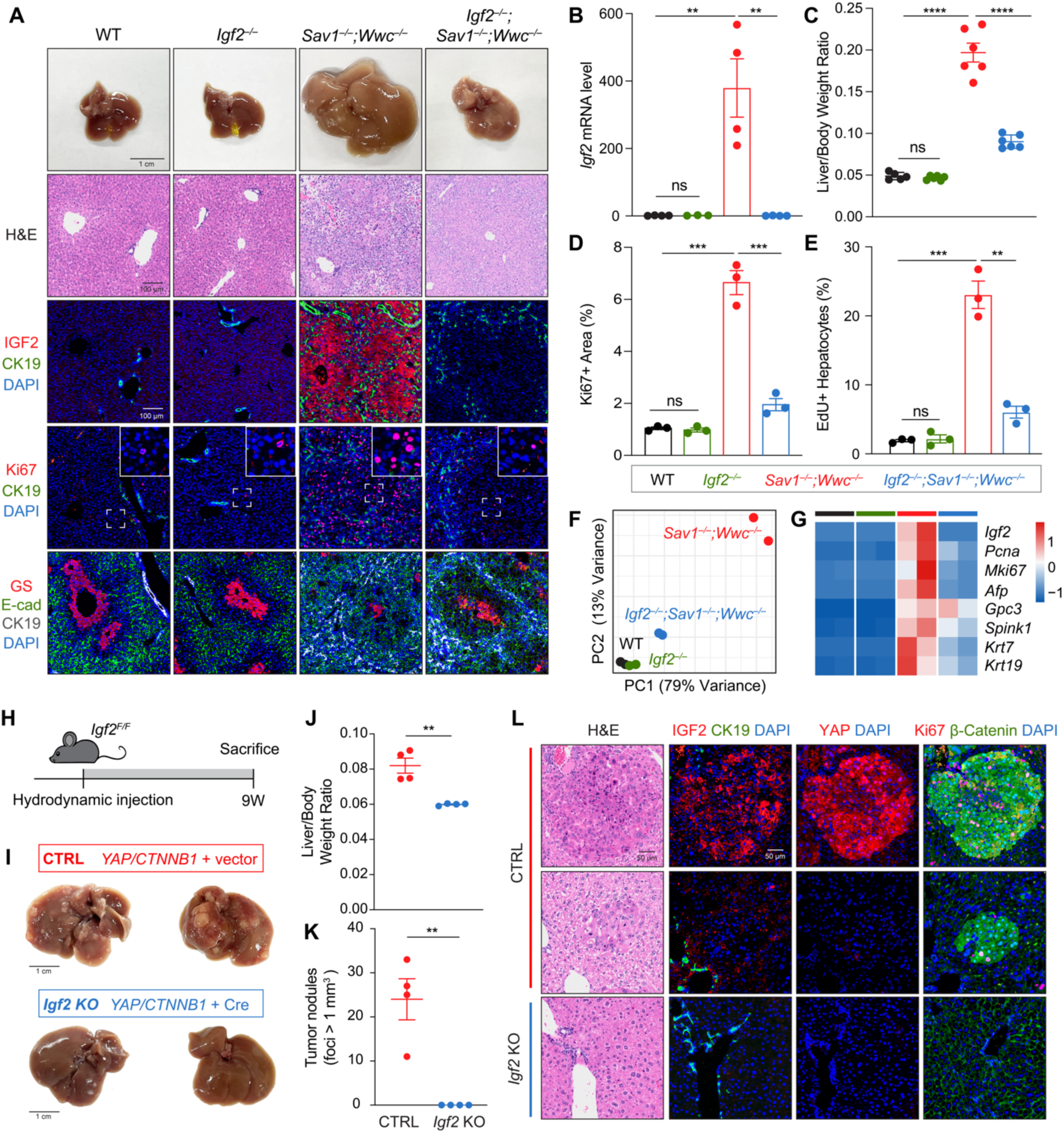
IGF2 is indispensable for hepatomegaly and tumorigenesis driven by YAP/TAZ activation. (A) *Igf2* deletion impairs YAP/TAZ-induced liver enlargement and cell proliferation. Gross liver images, H&E staining, and immunostaining of liver sections from 3-week-old WT, *Igf2^−/−^*, *Sav1^−/−^;Wwc*^−/−^, and *Igf2^−/−^;Sav1^−/−^;Wwc*^−/−^ mice are shown. E-cad: E-cadherin. Scale bars: 1 cm (gross liver image), 100 μm (H&E and immunostaining). (B) Quantitative RT-PCR results of *Igf2* expression. (C) Liver-to-body weight ratios of 3-week-old WT, *Igf2^−/−^*, *Sav1^−/−^;Wwc*^−/−^, and *Igf2^−/−^;Sav1^−/−^;Wwc*^−/−^ mice. (D and E) *Igf2* deletion impairs YAP/TAZ-induced cell proliferation. Quantifications of Ki67 (D) and EdU (E). EdU-positive hepatocytes were quantified by calculating the ratio of EdU-positive areas to HNF4α-positive areas; staining images are presented in Figure S5C. (F and G) Transcriptional profiles of 3-week-old livers. Principal component analysis (PCA) (F) and heatmap (G) are shown. (H) Schematic diagram illustrating the experimental procedure. *Igf2^F/F^* mice were subjected to hydrodynamic tail vein injections of YAP (*YAP-S127A*), β-Catenin (*CTNNB1-S33/37A*), and piggyBac (PB) to induce hepatocyte-specific gene expression. Cre plasmids were co-injected to delete *Igf2* in the *Igf2* knockout (KO) group (blue), while equal amounts of vector plasmids were injected in the control (CTRL) group (red). Livers were harvested nine weeks post-injection. (I-K) *Igf2* deletion significantly blocks HB development. Gross liver images (I), liver-to-body weight ratios (J), and tumor nodules (I) of the *Igf2* KO group (blue) and CTRL group (red) are shown. Scale bar in I: 1 cm. (L) H&E staining and immunostaining of liver sections from the CTRL and *Igf2* KO groups. Scale bar: 50 μm. Data in B-E, J, and K are presented as mean ± SEM, and each point represents an individual mouse. *P* values were assessed by unpaired Student’s t-tests. **p < 0.01, ***p < 0.001, ****p < 0.0001; ns: not significant.

YAP/TAZ regulates *IGF2* expression in HB (Figs 2H, I). We then asked whether YAP/TAZ-regulated *IGF2* expression is required for HB development. Using HDI and piggyBac (PB) transposon system, we delivered active YAP (*YAP-S127A*) and β-Catenin (*CTNNB1-S33/37A*), with or without a Cre recombinase, into the livers of *Igf2^F/F^* mice (Fig 5H). Expression of YAP and β-Catenin for 9 weeks resulted in robustly enlarged livers and visible tumor nodules (Fig 5I). Histological analysis revealed numerous dense, nearly spherical lesions with round nuclei, lacking severe necrosis, fibrosis, or inflammation, which are typical characteristics of epithelial HB (Fig 5L) ^13,46,53^. IGF2 was significantly upregulated in tumors with YAP expression (Fig 5L). However, it was undetectable in some small tumors expressing only β-Catenin (Fig 5L). In contrast, deletion of *Igf2* blocked HB development, resulting in relatively normal liver morphology and liver-to-body weight ratios (Figs 5I-K). Histological analysis did not detect tumor lesions or abnormal cell proliferation (Fig 5L). These results suggest that IGF2 deficiency impairs HB development induced by YAP/TAZ.

### YAP/TAZ-IGF2 induces IGF1R signaling in liver growth and tumorigenesis

To understand how IGF2 functions downstream of YAP/TAZ to regulate liver growth and hepatomegaly, we analyzed gene expression profiles of *Sav1^−/−^;Wwc*^−/−^ and *Igf2^−/−^;Sav1^−/−^;Wwc*^−/−^ livers. Consistent with attenuated cell proliferation observed in *Igf2*-deficient livers, the most significantly downregulated genes were involved in cell cycle and DNA replication. Conversely, the upregulated genes were primarily associated with metabolic functions such as bile acid secretion, gluconeogenesis, and lipid synthesis, indicating a restoration of normal liver function (Figs 6A and S5E).

**Figure 6.**
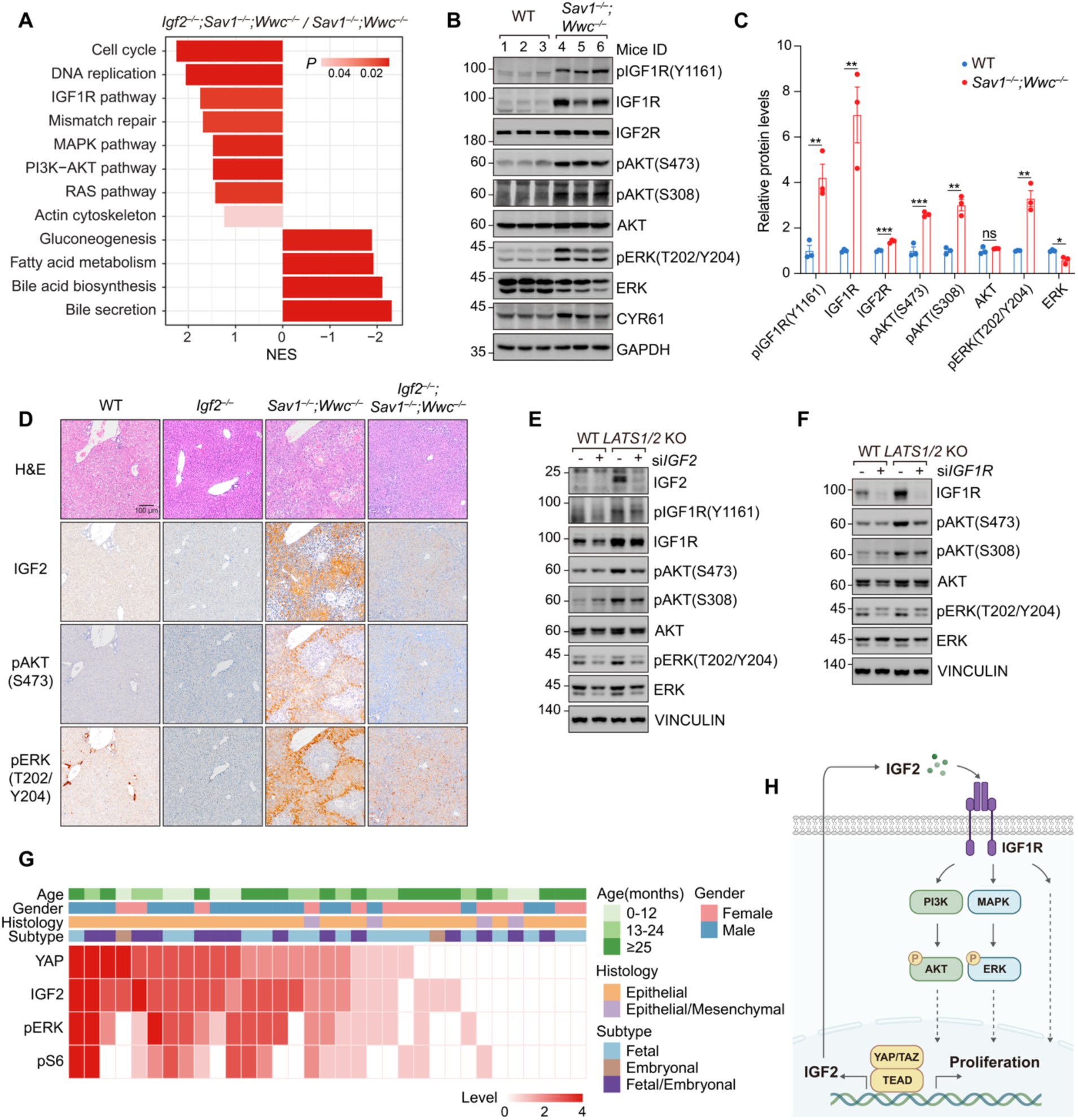
YAP/TAZ-IGF2 activates IGF1R signaling in hepatomegaly and tumorigenesis. (A) Gene Set Enrichment Analysis (GSEA) for the differentially expressed genes between *Igf2^−/−^;Sav1^−/−^;Wwc*^−/−^ and *Sav1^−/−^;Wwc*^−/−^ livers. (B and C) Activation of IGF1R-AKT/ERK signaling in *Sav1^−/−^;Wwc*^−/−^ livers. Protein samples were obtained from livers of 3-week-old mice and subjected to immunoblotting (B). Quantifications are presented as mean ± SEM, and each point represents an individual mouse (C). *P* values were assessed by unpaired Student’s t-tests. **p < 0.01, ***p < 0.001; ns: not significant. (D) H&E staining and immunostaining of IGF2, pAKT(S473), and pERK (T202/Y204) in liver sections from 3-week-old WT, *Igf2^−/−^*, *Sav1^−/−^;Wwc*^−/−^, and *Igf2^−/−^;Sav1^−/−^;Wwc*^−/−^ mice. Scale bar: 100 μm. (E and F) Knockdown of *IGF2* (E) or *IGF1R* (F) impairs AKT and ERK activation in *LATS1/2* knockout (KO) HEPG2 cells. (G) Phosphorylation of ERK and S6 is closely associated with IGF2 level and YAP activation in human HB specimens. (H) Schematic diagram illustrating that YAP/TAZ-induced IGF2 activates IGF1R signaling and promotes cell proliferation.

Notably, the deletion of *Igf2* significantly suppressed genes involved in the IGF1R, PI3K-AKT, and RAS-MAPK pathways (Figs 6A and S6A). IGF2 is a secreted factor that functions by binding to and activating specific cell surface receptors ^54^. As class II receptor tyrosine kinase, IGF1R is a functional receptor for IGF2, regulating cell proliferation, survival, and protein synthesis through activation of downstream pathways, including PI3K-AKT and RAS-MAPK ^55^. Interestingly, crosstalk between the Hippo pathway and PI3K signaling has been reported previously ^56,57^. We noticed that IGF1R, AKT, and MAPK were robustly activated in *Sav1^−/−^;Wwc*^−/−^ livers but attenuated in the absence of *Igf2*, as indicated by immunoblotting and immunostaining for phosphorylation of IGF1R, AKT, ERK, and S6 (Figs 6B-D and S6B). *In vitro*, *LATS1/2*-deficient HEPG2 cells also showed higher phosphorylation of IGF1R, AKT, and ERK, similar to the results observed in cells directly treated with recombinant IGF2 protein (Figs 6E, F and S6C). However, this activation was impaired upon knockdown of *IGF2* or *IGF1R* (Figs 6E, F). Hence, IGF2-IGF1R signaling plays a vital role in YAP/TAZ-induced AKT and MAPK activation during liver growth.

IGF1R signaling is closely associated with tumor malignancy ^54^. In IGF2-expressing liver tumors with dysregulated Hippo signaling, we detected increased phosphorylation of ERK and S6 (Fig S6D). Furthermore, in HB specimens, irrespective of age, gender, or pathological subtype, the phosphorylation of ERK and S6 closely correlated with *IGF2* expression and YAP activation (Figs 6G and S6E-G). These results indicate that YAP/TAZ-induced IGF2 activates IGF1R and downstream growth-promoting signaling during liver growth and tumor development (Fig 6H).

### Targeting IGF2-IGF1R signaling blocks YAP/TAZ-induced hepatomegaly and hepatoblastoma

Given that YAP/TAZ-induced IGF2 activates IGF1R signaling, we investigated whether targeting the IGF2-IGF1R axis could block hepatomegaly and tumorigenesis driven by dysregulated Hippo signaling. Picropodophyllin (PPP, also known as AXL1717) is a potent and selective IGF1R inhibitor with no overt effects on other structurally similar tyrosine kinase receptors, including the insulin receptor (IR) ^58^ (Fig S7A). We administered PPP or placebo daily to 10-day-old *WT* and *Sav1^−/−^;Wwc*^−/−^ mice and assessed liver size on postnatal day 21 (Fig 7A). Strikingly, IGF1R inhibition significantly suppressed liver enlargement and cell proliferation in *Sav1^−/−^;Wwc*^−/−^ livers (Figs 7B-D and S7B). However, PPP treatment in WT mice also reduced liver size compared to placebo treatment (Figs 7B-C and S7B), which was not observed in *Igf2^−/−^* livers (Figs 5A,C). In the presence of PPP, IGF2 secreted from the liver and other organs may fail to activate IGF1R and liver growth. Indeed, IGF1R inhibition blocked hepatomegaly induced by YAP/TAZ and exerted a growth-inhibitory effect on other organs, as evidenced by a gradual decrease in body weight (Fig S7C).

**Figure 7.**
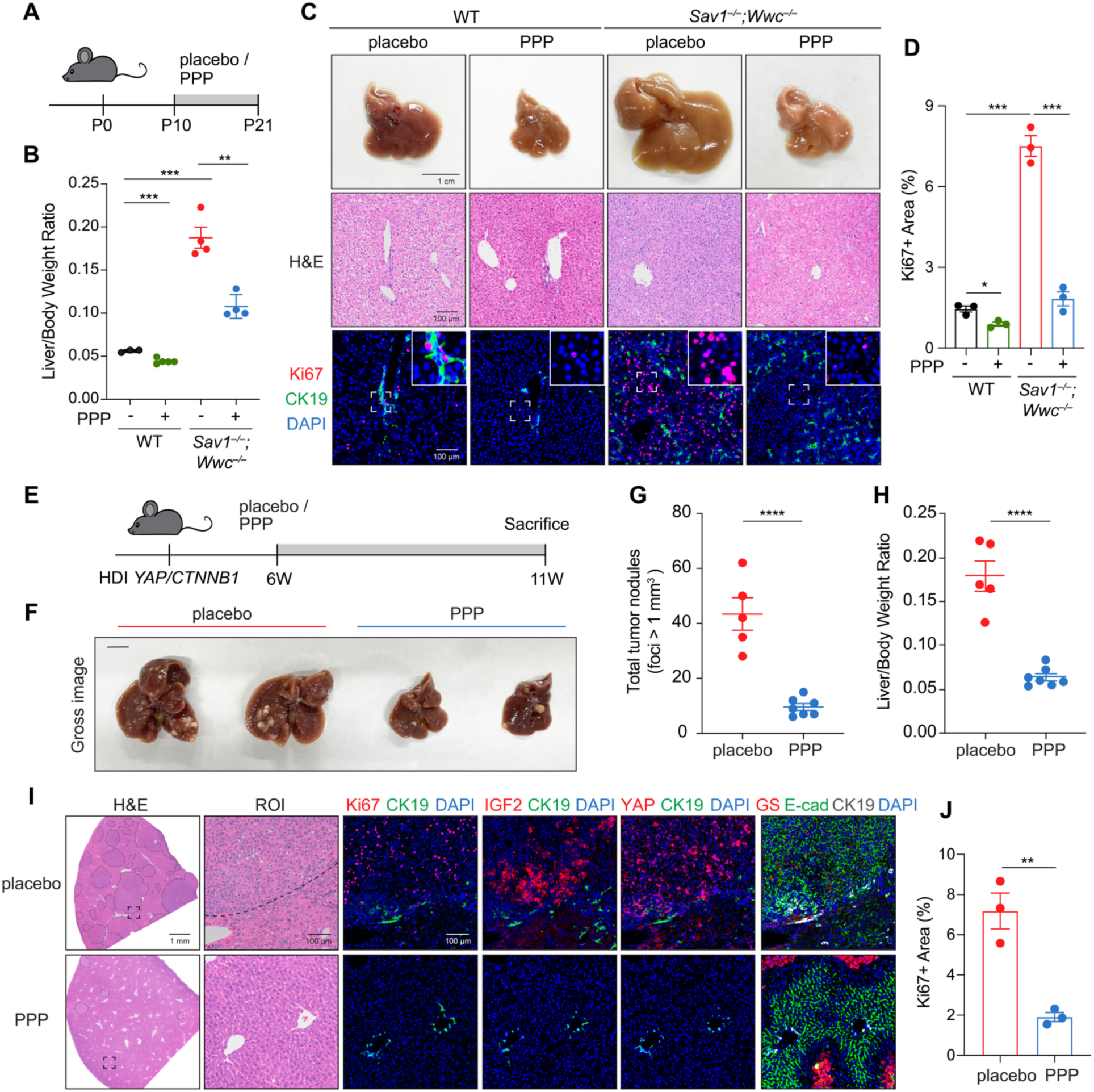
Targeting IGF2-IGF1R signaling blocks liver enlargement and hepatoblastoma driven by YAP/TAZ activation. (A) Schematic diagram illustrating the experimental procedure. WT and *Sav1^−/−^;Wwc*^−/−^ mice were administered intraperitoneally with either Picropodophyllin (PPP, 5 mg/kg) or placebo daily starting from postnatal day 10 (P10). Livers were harvested on P21. (B) Liver-to-body weight ratios of 3-week-old WT and *Sav1^−/−^;Wwc*^−/−^ mice, treated with either PPP or placebo. (C) PPP treatment suppresses cell proliferation and liver growth. Gross liver images, H&E staining, and immunostaining images of Ki67 (red), CK19 (green), and DAPI (blue). Scale bars: 1 cm (gross liver image), 100 μm (H&E and immunostaining). (D) Quantification results of Ki67 staining. (E) Schematic diagram illustrating the experimental procedure. Mice were subjected to hydrodynamic tail vein injections of YAP (*YAP-S127A*), β-Catenin (*CTNNB1-S33/37A*), and piggyBac (PB) to induce HB development. Six weeks after injection, mice were randomly divided into two groups and treated with either PPP (20 mg/kg) or placebo via intraperitoneal injection once daily. Livers were harvested after 11 weeks. (F-H) PPP treatment reduces tumor burdens. Gross liver images (F), total tumor nodules (G), and liver-to-body weight ratios (H). Scale bar in F: 1 cm. (I) H&E staining and immunostaining of liver sections from mice treated with placebo or PPP. Tumors are outlined with black dashed lines. E-cad: E-cadherin. Scale bars: 1 mm (H&E), 100 μm (ROI, immunostaining). (J) Quantification results of Ki67 staining. Data in B, D, G, H, and J are presented as mean ± SEM, and each point represents an individual mouse. *P* values were assessed by unpaired Student’s t-tests. *p < 0.05, **p < 0.01, ***p < 0.001, ****p < 0.0001.

HB is the most prevalent liver cancer in children, but effective and well-tolerated targeted therapies are currently unavailable ^53^. Aberrant activation of the YAP/TAZ-IGF2-IGF1R signaling axis in HB prompted us to test the effect of IGF1R inhibition on HB progression. In the *YAP/CTNNB1* mouse HB model, tumor lesions were observed 4 weeks post-HDI (Figs 7E and S7D). At 6 weeks post-HDI, we randomly assigned mice with HB into two groups: one treated daily with PPP and the other with placebo (Fig 7E). A lethal tumor burden appeared around 11 weeks post-HDI, accompanied by rapid weight loss in the placebo-treated mice (Fig S7E). However, mice receiving continuous PPP treatment showed no significant change in body weight or physical appearance (Fig S7E). Remarkably, while placebo-treated mice had enlarged livers with numerous HB lesions, PPP-treated mice exhibited smaller livers with fewer visible tumors (Figs 7F-H and S7F). Histological analysis confirmed a significant reduction in IGF2+, YAP+, and Ki67+ tumor cells following PPP treatment (Figs 7I, J and S7G). In addition, serological tests indicated a significant reduction in liver damage following PPP treatment, as evidenced by decreased ALT and AST levels (Figs S7H, I). These results underscore the potential of targeting IGF1R in treating HB driven by dysregulated Hippo signaling. Moreover, YAP/TAZ activation and IGF2 level may serve as molecular markers of tumors that likely respond to IGF1R inhibitors.

## DISCUSSION

The Hippo pathway is a growth control mechanism regulating organ size and tumorigenesis, but the key factor mediating its biological and pathological functions remains elusive. Our study demonstrates that the Hippo pathway directly regulates *IGF2* expression in the liver, specifically in hepatocytes. Through activating IGF1R signaling, IGF2 is indispensable for YAP/TAZ-induced hepatomegaly and HB in mice. Moreover, the Hippo-IGF2-IGF1R signaling axis can be potentially targeted for tissue regeneration and cancer treatment.

### Cryptic regulation of *IGF2* expression by the Hippo pathway

Tissue-specific transcription factors and epigenetic modifications are essential for maintaining cell identity and ensuring functional organ development ^59^. Unlike many other known target genes of the Hippo pathway, such as *CYR61* and *CTGF*, which are expressed in diverse cell types, YAP/TAZ induces IGF2 in hepatoblasts and hepatocytes, but not in cholangiocytes or liver mesenchymal cells. Furthermore, the activity of YAP/TAZ is deficient in matured hepatocytes, making the effect of *YAP/TAZ* deletion on *IGF2* expression undetectable in adult livers (Fig 8). Thus, IGF2 represents a cryptic target of YAP/TAZ, and this has not been uncovered in previous studies, likely due to the developmental stage- and cell type-specific regulations.

**Figure 8.**
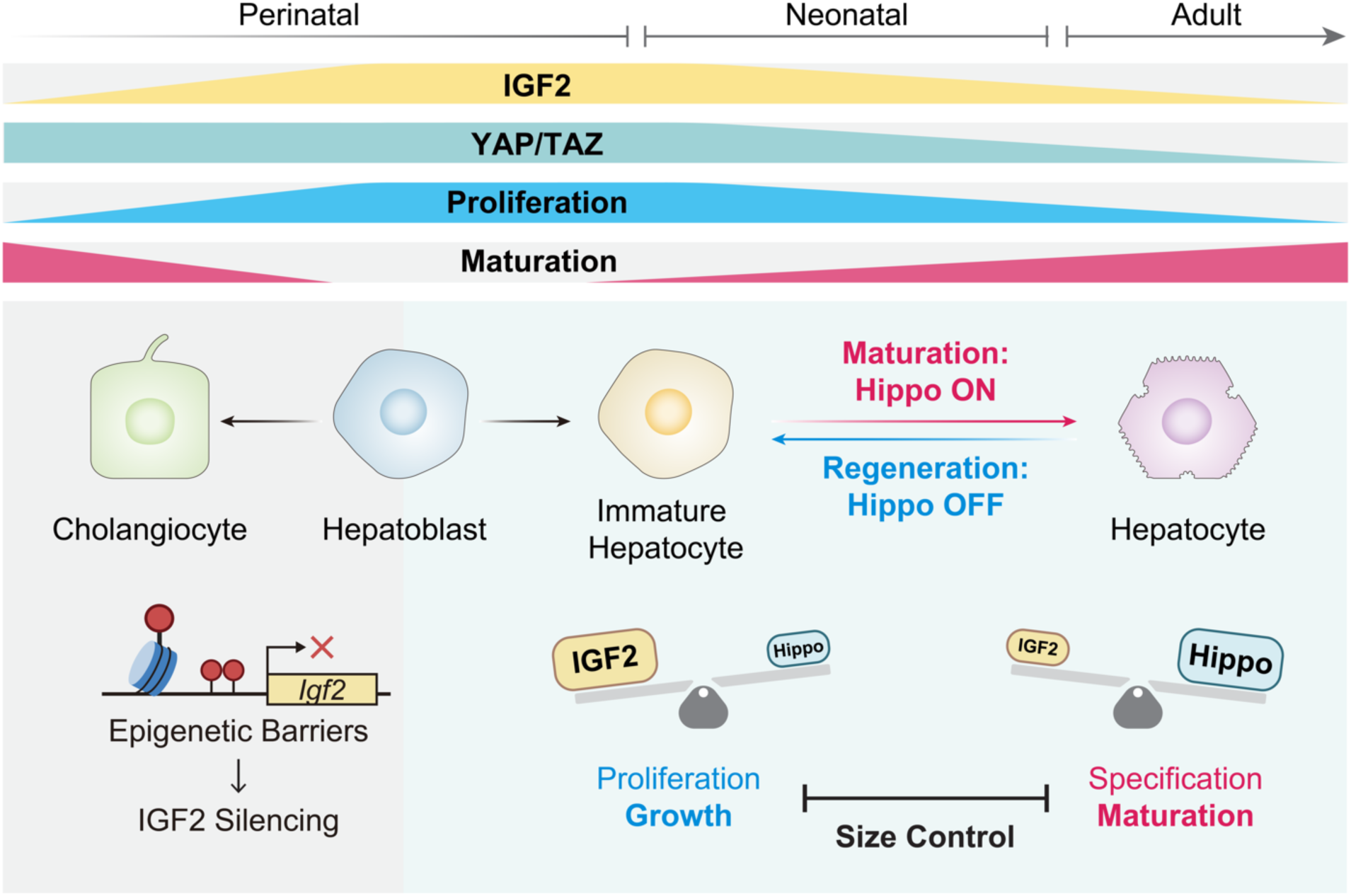
The regulation of IGF2 by the hippo pathway controls liver size. The dynamic expression of IGF2 and activity of the Hippo signaling (YAP/TAZ) intricately regulates liver growth and function. During liver development, hepatoblasts differentiate into immature hepatocytes and cholangiocytes. In immature hepatocytes, the gradual establishment of Hippo signaling (mainly HPO1) reduces YAP/TAZ activity, leading to *IGF2* silencing, attenuated hepatocyte proliferation, and enhanced hepatocyte maturation, which exerts a ceiling effect on liver size. Conversely, activation of YAP/TAZ in matured hepatocytes during liver regeneration induces their dedifferentiation into an immature state, resulting in *IGF2* reactivation, cell proliferation, and liver mass restoration. The Hippo-IGF2 axis determined whether hepatocytes assume an immature or matured state. The maturity of hepatocytes can instruct cells to undergo proliferation or specification, which controls liver growth and functional maturation, respectively. Hence, the antagonism between growth and maturation may ultimately control liver size. However, *IGF2* is not induced by YAP/TAZ in cholangiocytes due to epigenetic barriers, although YAP/TAZ activity is relatively high.

Differential epigenetic modifications in diverse cell lineages are responsible for the cell type-specific regulation of IGF2. Our findings reveal that DNA and histone methylation (mainly 5mC and H3K27me3) at *IGF2* promoters impedes its transcriptional activation by YAP/TAZ, likely by forming a non-permissive chromatin state that prevents TEAD binding (Figs 3, 4). In hepatocytes, *Igf2* fetal promoters are hypomethylated and adopt an open configuration, allowing gene expression to be effectively induced by YAP/TAZ. In contrast, other liver cells, particularly cholangiocytes, exhibit heavy H3K27me3 decoration at *Igf2* promoters, resulting in a closed state that prevents gene expression even when YAP/TAZ is active (Figs 4, 8). Differential H3K27me3 modifications may contribute to cell type-specific *IGF2* expression during liver development ^50,60^. In addition, the regulation of *IGF2* by YAP/TAZ is absent in many liver cancer cell lines, likely due to DNA methylation at the promoters and impaired TEAD binding ^33^. Other than 5mC and H3K27me3, additional mechanisms may also contribute to the cell type-specific expression of *IGF2*, which warrants further investigations.

The regulation of *IGF2* by the Hippo pathway is not limited to the liver. Intriguingly, in available transcriptome datasets, we have observed that *IGF2* expression is associated with YAP/TAZ activity in other cell types, including mouse embryonic stem cells, dorsal cranial neuroepithelium, and cardiac myocytes (Figs S8A-C) ^61–63^. In addition, in *Drosophila*, the expression of *insulin-like peptide 8* (*ILP8*) is regulated by Yorkie (the ortholog of YAP/TAZ) in a Scalloped (Sd, the ortholog of TEAD)-dependent manner (Fig S8D) ^64^. These findings suggest that the Hippo pathway can directly regulate *IGF2* expression in diverse organs and species.

### The Hippo-IGF2-IGFR signaling axis as a general organ size regulator

The IGF2-IGF1R signaling axis regulates organ growth in different species, including *Drosophila*, mice, and humans ^25,26,65–68^. Defective IGF2-IGF1R signaling affects cell proliferation and ultimately disturbs organ size and function ^29,30,52,69^. Thus, the activity of IGF1R needs to be spatiotemporally regulated during development and homeostasis. *IGF2* expression is strictly confined to fetal and neonatal stages and is silenced in adult organs ^27^. However, the molecular mechanism underlying *IGF2* silencing in adult tissues remains unclear. Our study suggests that the Hippo pathway is responsible for the postnatal silencing of *IGF2*, and the Hippo-IGF2-IGF1R signaling axis serves as a general mechanism for organ size control during development and regeneration (Fig 8).

The liver size is, in principle, mainly determined by the number of hepatocytes. The expression of *IGF2* during postnatal development is strongly associated with Hippo signaling and hepatocyte proliferation. The Hippo signaling in hepatocytes is gradually established after birth, which is inversely correlated with *IGF2* expression (Fig 8) ^10^. Reduced IGF2 downregulates IGF1R signaling and hepatocyte proliferation, which may exert a ceiling effect on liver growth and size. In contrast, moderate inactivation of Hippo signaling leads to the retention of hepatoblasts or the dedifferentiation of hepatocytes into an immature state. In immature hepatocytes, *IGF2* expression is induced by YAP/TAZ, promoting cell proliferation and liver enlargement. Therefore, the Hippo-IGF2-IGF1R signaling axis is precisely regulated spatiotemporally to ensure proper liver development. An antagonism between the proliferation and specification of hepatocytes, or the growth and maturation of the liver, may determine the final organ size and functionality (Fig 8).

After damage or surgical resection, the restoration of liver size and function relies on effective regenerative mechanisms frequently associated with elevated YAP/TAZ activity ^70–72^. Transient YAP/TAZ activation has been reported following liver damage, and their deletion significantly compromises liver regeneration ^71,73,74^. IGF2 is also upregulated after liver damage, driving hepatocyte proliferation during regeneration ^75,76^. Although *IGF2* expression is silenced in adult hepatocytes, its fetal promoters remain permissive and can be reactivated by YAP/TAZ. Indeed, in the livers of aged mice, YAP/TAZ activation induces robust *IGF2* expression and liver growth (Figs 1G-J and S1B-D). Hence, the Hippo-IGF2-IGF1R signaling axis may be awakened by damage signals to promote liver regeneration (Fig 8). Similar mechanisms may operate in additional organs and could be harnessed for regenerative medicine.

### IGF2-IGF1R as a molecular target of cancers driven by YAP/TAZ

Overexpression of *IGF2* occurs in diverse cancers, including aggressive HB, glioblastoma, ependymoma, Ewing’s sarcoma, and rhabdomyosarcoma ^32–34,77,78^. In these tumors, sustained activation of IGF2-IGF1R signaling is considered a crucial oncogenic mechanism, as downstream AKT and ERK promote cancer cell proliferation and resistance to chemotherapy ^32,34,79–81^. Moreover, IGF2 may promote tumor growth by remodeling the tumor microenvironment ^82–84^. Consequently, tumors with high *IGF2* expression are typically associated with poor prognosis, characterized by rapid progression, frequent recurrence, and high mortality ^35,36,80^. Intriguingly, in addition to HB, the expression of *IGF2* is highly correlated with canonical YAP/TAZ target genes in many cancers from different tissue origins, such as breast, colon, esophagus, pancreas, prostate, and stomach, suggesting that YAP/TAZ regulates *IGF2* expression in multiple cancers (Fig S8E-J).

The Hippo pathway is one of the major oncogenic signaling pathways ^3,15,85^. However, therapeutic approaches targeting the Hippo pathway are somewhat limited, with recent developments mainly focusing on blocking the interaction between YAP/TAZ and TEAD ^15,86,87^. Deletion of *Igf2* or inhibition of IGF1R significantly blocks the progression of HB in mice, indicating that IGF2-IGF1R signaling is essential for YAP/TAZ-driven tumorigenesis (Figs 5 and 7). Therefore, the IGF2-IGF1R signaling axis represents a promising molecular target for treating cancers driven by YAP/TAZ. Several IGF1R inhibitors and IGF2-neutralizing antibodies are currently in clinical trials for cancer therapies ^88,89^. It would be attractive to evaluate whether these modalities are effective in treating cancers with a YAP/TAZ dependency.

## Supporting information

Resource Table

## Acknowledgments

We thank Dr. Cun Wang for providing the SNU475 cell line. This study is supported by grants from the National Key R&D Program of China (2020YFA0803202, 2018YFA0800304), the National Natural Science Foundation of China (32425017, 32370770, 32200570, 82403053), the Science and Technology Commission of Shanghai Municipality (21S11905000), and Shanghai Municipal Health Commission (2022XD049). This work is also supported by the Medical Science Data Center and the Core Facility at Shanghai Medical College of Fudan University.

## Data and material availability

The RNA-seq (CRA019258), scRNA-seq (CRA016280), WGBS (CRA019266), targeted methylation sequencing (HRA008706), and ChIP-seq (CRA019350) data generated in this study have been deposited in the Genome Sequence Archive in National Genomics Data Center, China National Center for Bioinformation/Beijing Institute of Genomics, Chinese Academy of Sciences (PRJCA030536), and are publicly accessible at https://ngdc.cncb.ac.cn/gsa.

## EXPERIMENTAL MODEL AND SUBJECT DETAILS

### Genetic mouse models

All mouse experiments were approved by the Animal Ethics Committee of Shanghai Medical College, Fudan University, and conducted in compliance with institutional guidelines. Male and female mice were randomly assigned to experiments. *Igf2^F/F^* mice were generated as described in Figure S5A. *Wwc1^F/F^*, *Wwc2^F/F^*, and *Nf2^F/F^* mice were generated in our previous work ^9,90^. *Sav1^F/F^*, *Mst1^F/F^*, *Mst2^△/△^*, *Lats1^F/F^*, *Lats2^F/F^*, *Yap^F/F^*, and *Taz^F/F^* mice were described previously ^91–95^. Aside from *Mst1^F/F^;Mst2^△/△^* mice, which were on a mixed background of C57BL/6J and 129/SvJ, all other mice used in this study were on a C57BL/6J background. *Albumin-Cre* (*Alb-Cre*, The Jackson Laboratory, #003574) mice were used to delete genes in the liver during development. For liver-specific acute gene deletion, Ad-*Cre* (2×10^9^ pfu/mouse, HANBIO, #HBAD-1010) or AAV8-*TBG-Cre* (1×10^11^ vg/mouse, Obio Technology, #H5721) was injected into mice via the lateral tail vein. *Rosa26-tdTomato* reporter mice (Ai9, The Jackson Laboratory, #007905), which express robust tdTomato upon Cre-mediated recombination, were used to assess Cre efficiency and trace lineage. At the indicated age, mouse livers were harvested and either fixed in formalin solution and paraffin-embedded, or stored in liquid nitrogen for subsequent gene expression analysis.

### Mouse cancer models

Hepatoblastoma (HB), hepatocellular carcinoma (HCC), and intrahepatic cholangiocarcinoma (ICC) models were established using the hydrodynamic injection (HDI) method with piggyBac (PB) and Sleeping Beauty (SB) transposase systems. For the *YAP/CTNNB1* model, plasmids containing 20 μg YAP(S127A), 20 μg CTNNB1(S33/37A), 28 μg PB, and 100 μg Cre (or empty vector) were diluted in 2 mL 0.9% saline, filtered, and injected into the lateral tail vein of 6-week-old mice at a volume equal to 10% body weight within 5-7 seconds. For the *CTNNB1/RAS* model, each mouse received injections of 25 μg CTNNB1(S33/37A), 25 μg NRAS(G12V), and 10 μg PB. Similarly, for the *AKT/RAS* model, 25 μg myr-AKT, 25 μg NRAS(G12V), and 10 μg PB were delivered to each mouse. For the *NICD/AKT* model, each mouse was injected with 25 μg NICD, 25 μg myr-AKT, and 10 μg SB. In the DEN-induced HCC model, 2-week-old mice were given a single intraperitoneal injection of diethylnitrosamine (DEN, 25 mg/kg, Sigma-Aldrich, #N0756) and harvested approximately 10 months later.

### Cell lines and cell culture

All cell lines were grown in tissue culture incubators at 37°C with 5% CO_2_. Human HB (HEPG2 and HUH6), HCC (HUH7, SNU449, and SNU475), HEK293A, and HEK293T cell lines were cultured in DMEM medium supplemented with 10% fetal bovine serum (FBS). Human ICC (RBE and SG231) cell lines were cultured in RPMI-1640 medium with 10% FBS. Primary mouse hepatocytes were plated on type I collagen-coated plates and cultured in William’s E medium with 10% FBS, 0.05% insulin, and 1 mM sodium pyruvate. *LATS1/2* knockout HepG2 cells were generated in this study using CRISPR/*Cas9* system, with gene deletion confirmed via immunoblotting. Stable cell lines expressing YAP, YAP^2SA^ (YAP-S127/397A), YAP^2SA,S94A^ (YAP-S94/127/397A), TAZ^4SA^ (TAZ-S66/89/117/311A), LATS1, and LATS2 were established through lentiviral transduction.

### Human hepatoblastoma specimens

Human hepatoblastoma specimens for immunostaining analysis were collected from the Department of Pediatric Surgery at Children’s Hospital of Fudan University. Informed consent was obtained from all patients prior to sample collection, and corresponding clinical information is provided in Table S1. The use of these specimens for research was approved by the Ethics Committee of Shanghai Medical College, Fudan University.

## METHOD DETAILS

### Plasmid construction, lentivirus production, and transduction

Gene cloning was performed using PrimeSTAR Max DNA Polymerase (Takara, #R045A), and gene fragments were inserted into the pLVX-puro or pLVX-hygro vector with a FLAG tag via the ClonExpress MultiS One Step Cloning Kit (Vazyme, #C113). For lentivirus production, HEK293T cells were co-transfected with pLVX-based plasmids, psPAX2 (Addgene, #12260), and pMD2.G (Addgene, #12259) packaging vectors at a ratio of 4:3:1 using polyethyleneimine (PEI, Polysciences, #23966-2). Virus-containing medium was collected at 48 and 72 hours post-transfection, filtered, and centrifuged at 32,000 rpm for 2 hours at 4 °C. The viral pellets were resuspended in DMEM medium and used for transduction in the presence of Polybrene (1:1000, Sigma-Aldrich, #TR-1003). After 48 hours, transduced cells were selected in medium containing puromycin (1 μg/mL, InvivoGen, #ant-pr-1) or hygromycin B (100 μg/mL, InvivoGen, #ant-hg-1).

### CRISPR/Cas9-mediated gene knockout

Gene-specific single-guide RNA (sgRNA) sequences were designed using the E-CRISP website (http://www.e-crisp.org/E-CRISP/). These sgRNAs were cloned into the lentiCRISPR v2 vector (Addgene, #52961) and co-transfected with psPAX2 and pMD2.G into HEK293T cells for lentivirus production. Targeted cells were transduced with the lentivirus and selected in medium containing puromycin (1 μg/mL, InvivoGen, #ant-pr-1) for 2-3 days. Gene knockout efficiency was validated by immunoblotting. The sgRNA sequences are listed in Table S2.

### Isolation of mouse hepatocytes

Primary mouse hepatocytes were isolated using a modified two-step collagenase perfusion method ^96^. Following anesthesia and abdominal incision, a catheter was inserted into the inferior vena cava, and Hanks’ buffer I (Sigma-Aldrich, #H2387) was perfused at a flow rate of 10 mL/min. Once the liver became pale, the portal vein was quickly cut, and the superior vena cava was clamped. After perfusion with Hanks’ buffer I for approximately 5 minutes, pre-warmed digestion buffer containing type IV collagenase (Sigma-Aldrich, #C5138) and Ca^2+^ was infused at a rate of 3-4 mL/min. When the liver developed cracks and became gelatinous, it was minced and passed through a 200 μm filter. Isolated hepatocytes were washed twice with Hanks’ buffer II (Sigma-Aldrich, #H6136), resuspended in William’s E medium (Sigma-Aldrich, #W4125), and used for culture, crosslinking, or nucleic acid extraction.

### RNA interference

Cells were plated in 12-well plates with 500 μL DMEM medium and cultured for 12 hours before transfection. For transfection, 12 pmol small interfering RNA (siRNA) and 2 μL Lipofectamine RNAiMAX Transfection Reagent (Invitrogen, #13778075) were each diluted in 50 μL serum-free medium, mixed gently, and incubated for 5 minutes at room temperature before being added to the cells. Cells were incubated for 48-72 hours post-transfection before analysis. The siRNA sequences were designed using the BLOCK-iT RNAi Designer (https://rnaidesigner.thermofisher.com/rnaiexpress/) and are listed in Table S3.

### RNA extraction, reverse transcription, and quantitative real-time PCR

Total RNA was extracted using the TaKaRa MiniBEST Universal RNA Extraction Kit (Takara, #9767) according to the manufacturer’s instructions. For each sample, 1 μg RNA was reverse transcribed into cDNA using the TransScript One-Step gDNA Removal and cDNA Synthesis SuperMix (TransGen Biotech, #AT311). Quantitative real-time PCR (qPCR) was performed using the TB Green Premix Ex Taq (Takara, #RR420A) on the QuantStudio 6 Flex Real-Time PCR System (Applied Biosystems). All qPCR experiments were conducted with at least three biological replicates, and relative quantification was calculated using the 2^−ΔΔCT^ method. The primers used for qPCR are listed in Table S4.

### Bulk RNA sequencing and analysis

RNA was extracted using the TaKaRa MiniBEST Universal RNA Extraction Kit (Takara, #9767). For library preparation, total RNA (1 μg per sample) was processed with the VAHTS mRNA-seq V10 Library Prep Kit for Illumina (Vazyme, #NR606) following the manufacturer’s instructions. Libraries were sequenced on the NovaSeq platform (Illumina) using a 2 × 150 bp paired-end run. Sequenced reads were aligned using the spliced read aligner TopHat (v2.1.1) with the GRCm38 (mm10) mouse genome or GRCh38 (hg38) human genome as reference genomes. BigWig files were generated with deeptools (v3.1.3) and visualized using Integrative Genomics Viewer (IGV, v2.16.0). Gene expression levels were estimated with cufflinks (v2.2.1) and featureCounts (v2.0.6), and normalized by transcripts per million (TPM). Differential expression analysis was performed using the R package DESeq2 (v1.38.3). Principal component analysis (PCA) was conducted with the R package gmodels (v2.19.1). Gene Set Enrichment Analysis (GSEA) was run on the ranked list using the Molecular Signatures Database (MSigDB) as the gene sets. YAP signature scores were calculated using the Cordenonsi Hippo signature gene set ^97^. Visualizations were generated using R packages ggplot2 (v3.5.1), ggrepel (v0.9.5), and pheatmap (v1.0.12). The signature gene sets used in this study are listed in Table S6. Sample information and TPM values are provided in Table S7 and S8.

### Single-cell RNA sequencing and analysis

Mouse livers were surgically removed and kept in MACS Tissue Storage Solution (Miltenyi Biotec, #130-100-008) until processing. After washing with PBS, livers were minced into 1 mm^3^ pieces and digested with 1 mg/mL dispase (Sigma-Aldrich, #D4818), 1.5 mg/mL type II collagenase (Sigma-Aldrich, #C2-22), and 30 U/mL DNase I (Sigma-Aldrich, #DN25) for 20 minutes at 37°C with agitation. Digested livers were filtered through a 70 μm filter, centrifuged, and suspended in red blood cell lysis buffer (Miltenyi Biotec, #130-094-183). After washing with PBS, cells were resuspended in PBS containing 0.04% BSA, filtered again through a 40 μm filter, and assessed for viability using a Fluorescence Cell Analyzer (Countstar). Libraries were generated using the Chromium Single Cell 3’ V3.1 Reagent Kit (10X Genomics) and Chromium Controller Instrument (10× Genomics). Briefly, cells were loaded into each channel to generate single-cell Gel Bead-In-Emulsions (GEMs). After reverse transcription, GEMs were broken, and barcoded cDNA was purified, amplified, fragmented, and ligated with adaptors. The libraries were quantified using a Qubit 4 fluorometer (Invitrogen), and their size distribution was determined using a Bioanalyzer 2200 (Agilent). Libraries were sequenced on the NovaSeq platform (Illumina) using a 2 × 150 bp paired-end run.

For data analysis, sequenced reads were filtered using the fastp pipeline ^98^ and aligned to the GRCm38 (mm10) mouse genome using CellRanger (v6.6.1). After removing outliers and doublets based on library size, number of expressed genes, and mitochondrial gene content, the data were further processed for normalization, scaling, identification of variable features, and PCA calculation using the R package Seurat (v4.3.0.1). The Seurat RunUMAP function was used to perform dimensionality reduction and unsupervised clustering. The Wilcoxon rank-sum test was used to identify markers in each cluster, and cell types were assigned based on these markers. Signature scores for selected gene sets were calculated using the Seurat AddModuleScore function. Visualizations were generated using R packages ggplot2 (v3.5.1) and Seurat (v4.3.0.1).

### Immunoblotting

Cells were lysed directly in 1× SDS loading buffer (50 mM Tris-HCl at pH 6.8, 2% SDS, 10% glycerol, 0.025% bromophenol blue, and 1% β-mercaptoethanol). Tissue samples were lysed in RIPA buffer (50 mM HEPES at pH 7.5, 150 mM NaCl, 0.1% SDS, 0.5% sodium deoxycholate, and 1% Triton X-100) supplemented with freshly added protease and phosphatase inhibitors, followed by the addition of 4× SDS loading buffer. Proteins were separated by SDS-PAGE, and transferred to nitrocellulose membranes. Membranes were blocked with 5% non-fat milk for 1 hour at room temperature, incubated overnight at 4°C with primary antibodies diluted in 5% bovine serum albumin (BSA), and then incubated with HRP-conjugated secondary antibodies diluted in 5% milk for 1 hour at room temperature. Detection was performed using the High-sig ECL Immunoblotting Substrate (Tanon, #180-501), and chemiluminescence was captured using the 5200S chemiluminescent imaging system (Tanon). Primary antibodies used in immunoblotting are listed in Table S5.

### Immunofluorescence

For cultured cells, coverslips were fixed in 4% paraformaldehyde for 10 minutes, washed three times with PBS, and permeabilized with 0.2% Triton X-100 in PBS for 5 minutes. For paraffin-embedded tissues, 5 μm sections were rehydrated, subjected to heat-induced antigen retrieval using 10 mM sodium citrate buffer for 20 minutes, and washed three times with PBST (PBS containing 0.1% Triton X-100). After blocking with 10% goat serum in PBST for 1 hour, coverslips or sections were incubated overnight at 4°C with primary antibodies diluted in PBST containing 1% BSA. Samples were then incubated with Alexa Fluor 488-, Alexa Fluor 555-, or Alexa Fluor 647-conjugated secondary antibodies and DAPI at room temperature for 1 hour, followed by mounting with Fluoromount-G (Invitrogen, #0100-01). Images were captured using an LSM900 confocal microscope (Zeiss) and a VS200 Slide Scanner (Olympus). Primary antibodies used in immunofluorescence are listed in Table S5.

### Immunohistochemistry

Paraffin-embedded tissues were sectioned, rehydrated, and underwent heat-induced antigen retrieval using 10 mM sodium citrate buffer for 20 minutes. Tissue sections were treated with 3% H_2_O_2_ for 30 minutes to quench endogenous peroxidase activity, followed by blocking with 3% BSA in TBST (TBS containing 0.1% Triton X-100) for 1 hour at room temperature. After overnight incubation at 4°C with primary antibodies, sections were washed three times with TBST and incubated with HRP-conjugated secondary antibodies (DAKO, #K4002). Visualization was performed using the DAB detection kit (GeneTech, #GK500705). Sections were then counterstained with hematoxylin, dehydrated, and mounted using mounting medium (BaSo, #BA7004). Images were captured using an Axioscope 5 Microscope (Zeiss) or a VS200 Slide Scanner (Olympus). Primary antibodies used in immunohistochemistry are listed in Table S5.

### *In situ* EdU labeling

For *in situ* cell proliferation analysis in mouse livers, 50 mg/kg EdU (Carbosynth, #NE08701) was injected intraperitoneally 24 hours before sacrifice. Liver sections were hydrated and underwent antigen retrieval. EdU staining was performed using the Cell-Light Apollo567 Stain Kit (Ribobio, #C10371-1) according to the manufacturer’s instructions. Sections were then incubated overnight with anti-HNF4α (CST, #3113, 1:100 diluted) and anti-CK19 (Abcam, #ab52625, 1:200 diluted) primary antibodies, followed by incubation with secondary antibodies and mounting as described above. Images were captured using an LSM900 confocal microscope (Zeiss) and a VS200 Slide Scanner (Olympus). The EdU- and HNF4α-positive areas were measured and normalized to assess hepatocyte proliferation using ImageJ (NIH).

### Targeted methylation sequencing

Genomic DNA was extracted using the DNeasy DNA Isolation Kit (QIAGEN, #69504) and underwent bisulfite conversion using the EZ DNA Methylation-Gold Kit (ZYMO, #D5005) according to the manufacturer’s instructions. The converted DNA samples were amplified by PCR targeting specific genomic regions, followed by adaptor ligation with index sequences. DNA libraries were sequenced on the NovaSeq platform (Illumina) using a 2 × 150 bp paired-end run. Filtered R1 and R2 sequencing reads were merged using FLASH (v1.2.11), and the merged reads were aligned to the reference genome using BLAST+ (v2.7.1). DNA 5-methylcytosine (5mC) levels at specific CpG sites were calculated using the formula: DNA 5mC level = Number of reads with methylation at the CpG site / Total number of reads at the CpG site. Visualizations were generated using the R package ggplot2 (v3.5.1). Primers used for PCR amplification are listed in Table S4. The DNA 5mC levels at *IGF2 P3/P4* region are provided in Table S9.

### Whole Genome Bisulfite Sequencing

Genomic DNA was extracted using the DNeasy DNA Isolation Kit (QIAGEN, #69504). For each sample, 1 μg DNA was randomly sheared to fragments less than 500 bp using an S220 ultrasonicator (Covaris), followed by end repairing and adapter ligation. After size selection of adapter-ligated fragments, a proportion of lambda DNA (negative control) was spiked in. Unmethylated cytosines were converted to uracil via bisulfite conversion, and sequencing was performed on the NovaSeq platform (Illumina). Sequenced reads were trimmed using Cutadapt (v1.9.1), aligned to the GRCm38 (mm10) mouse genome. Methylated cytosine (mC) calling was performed using Bismark (v0.7.12). BigWig files were generated using deeptools (v3.1.3) and visualized with Integrative Genomics Viewer (IGV, v2.16.0). Methylation levels at specific CpG sites were visualized using the R package ggplot2 (v3.5.1).

### Chromatin immunoprecipitation, sequencing and analysis

Cells were crosslinked with 1% formaldehyde for 10 minutes at room temperature and quenched with 0.125M glycine. Crosslinked cells were washed with ice-cold PBS, resuspended in lysis buffer (50 mM Tris-HCl at pH 7.4, 500 mM NaCl, 2 mM EDTA, 1% Triton X-100, 0.1% SDS, and 0.1% sodium deoxycholate) containing freshly added protease and phosphatase inhibitors, and sonicated using a Q800R3 Sonicator (Qsonica). The sheared chromatin was incubated overnight at 4°C with specific antibodies, and then immobilized on protein A/G beads (Smart-lifesciences, #SM015) for another 2 hours. For TEAD4, the chromatin was diluted to 0.05% SDS before incubation and then immobilized on 20ul Dynabeads Protein A (Thermo Fisher Scientific, #10002D) and 20ul Dynabeads Protein G (Thermo Fisher Scientific, #10004D). The bound fractions were washed three times with lysis buffer, twice with low-salt buffer (10 mM Tris-HCl at pH 8.0, 250 mM LiCl, 1 mM EDTA, 0.5% NP-40, and 0.5% sodium deoxycholate), and once with TE buffer (10 mM Tris-HCl at pH 8.0 and 1 mM EDTA). Elution and reverse crosslinking were performed in elution buffer (50 mM Tris-HCl 8.0, 10 mM EDTA, and 1% SDS) at 65°C for 5 hours with RNase A and proteinase K. DNA samples were purified using the PCR Purification Kit (Qiagen, #28104). For ChIP-qPCR, ChIP signals were analyzed using specific primers (listed in table S4) and normalized to input DNA. For ChIP-seq, 750 ng spike-in *Drosophila* chromatin and spike-in antibody (Active Motif, #61686) were added to each reaction before overnight incubation. DNA libraries were constructed using the VAHTS Universal DNA Library Prep Kit for Illumina (Vazyme, #ND607) and sequenced on the NovaSeq platform (Illumina). Sequenced reads were trimmed by trim_galore (v0.6.10) and mapped to the GRCm38 (mm10) mouse genome and dm6 *Drosophila* genome using bowtie2 (v2.2.5). Aligned results were normalized with spike-in reads. BigWig files were generated using deeptools (v3.1.3) and visualized with Integrative Genomics Viewer (IGV, v2.16.0). Primary antibodies used in ChIP are listed in Table S5.

### Epigenetic inhibitor treatment

Cells were plated and cultured for at least 12 hours before treatment. Decitabine (Selleck, #S1200, 50 μM), EPZ6438 (Selleck, #S7128, 5 μM), GSK343 (Selleck, #S7164, 5 μM), CPI-455 (Selleck #S8287, 5 μM), UNC0642 (Selleck, #S7230, 5 μM), CHIR99021 (Sigma, #SML1046, 5 μM), TSA(MCE, #HY-15144, 5 μM), SAHA(Selleck, #S1047, 5 μM), or ORY-1001(MCE, #HY-12782T, 5 μM) was added to the culture medium, with fresh drug-containing medium replaced after 48 hours. After 96 hours of treatment, genomic DNA, RNA, and proteins were extracted as described above.

### Picropodophyllin treatment

To investigate the effect of IGF2-IGF1R inhibition on liver size during development, 5 mg/kg Picropodophyllin (PPP, Selleck, #S7668) or placebo/vehicle (5% DMSO, 40% PEG300, 5% Tween80, 50% H_2_O) was administered intraperitoneally to mice once daily, starting from postnatal day 10, with analysis conducted on day 21. In YAP-driven HB models, plasmids containing 20 μg YAP^S127A^, 20 μg CTNNB1, and 8 μg PB were diluted in 2 mL 0.9% saline and injected via the lateral tail vein into 6-week-old mice in a volume equal to 10% of their body weight. Six weeks after injection, mice were randomly divided into two groups and treated with either 20 mg/kg Picropodophyllin or placebo/vehicle via intraperitoneal injection once daily. Body weight and overall condition were continuously monitored, and the livers were harvested after 11 weeks.

### ALT and AST measurement

Alanine aminotransferase (ALT) and aspartate aminotransferase (AST) levels in mouse sera were measured using a CA-200 automatic biochemical analyzer (URIT Medical Electronic), following the manufacturer’s guidelines.

### Publicly available sequencing data and analysis

The RNA-seq data from *YAPTg* livers (GSE178227) ^22^, mouse livers with *Nf2*, *Sav1*, *Wwc1/2*, or compound deletions (CRA008364) ^10^, mouse embryonic stem cells (GSE157706) ^62^, dorsal cranial neuroepithelium (GSE182721) ^61^, cardiac myocytes (GSE176142) ^63^, *Drosophila* eyes (GSE211458) ^64^, and the scRNA-seq data from developing mouse livers (GSE171993) ^23^ were analyzed as described above. The MBD-seq data from *YAPTg* livers (GSE178227) were analyzed as previously described ^22^. Processed snATAC-seq data from mouse livers were obtained from the UCSC Genome Browser (https://genome.ucsc.edu/s/cbravo/Bravo_et_al_Liver) ^51^. DNA methylation and gene expression data from various liver cancer cell lines were accessed from the CCLE datasets of the Cancer Dependency Map Project (Depmap, https://depmap.org/portal/) ^99^. Gene expression and Spearman correlation analysis of The Cancer Genome Atlas (TCGA) data were conducted through the UCSC Xena platform (https://xenabrowser.net).

## QUANTIFICATION AND STATISTICAL ANALYSIS

Statistical analyses were performed using R (v4.2.3) and GraphPad Prism (v9, GraphPad Software). Experiments were repeated at least three times, with results presented as mean ± SEM, as described in the figures and corresponding legends. Statistical significance was determined using unpaired Student’s t-tests or one-way analysis of variance (ANOVA). Correlation coefficients were analyzed using Spearman’s correlation. Statistical significance was denoted as follows: *p < 0.05, **p < 0.01, ***p < 0.001, ****p < 0.0001; ns indicates not significant.

---

**Figure S1.**
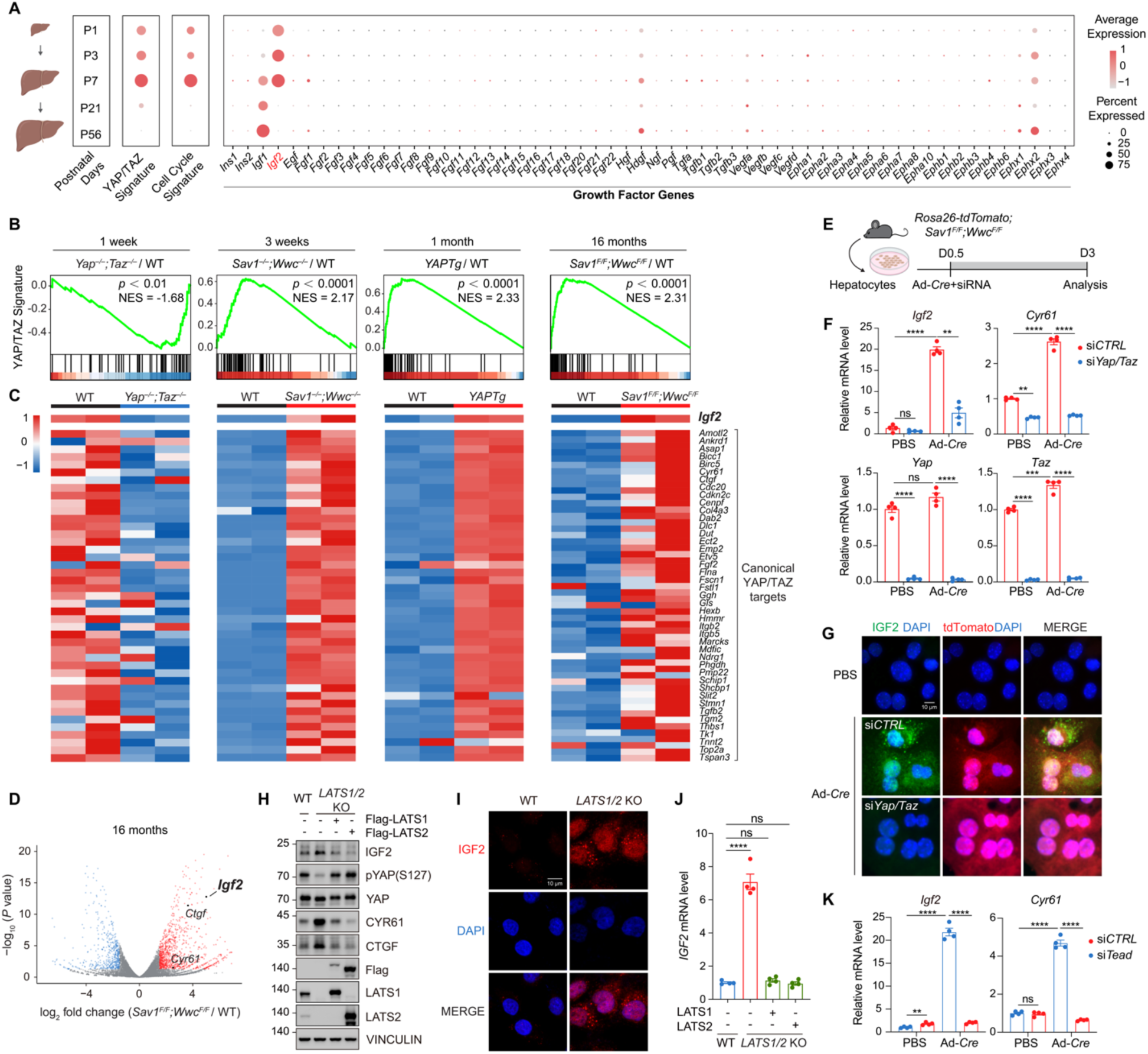
YAP/TAZ activation induces *IGF2* expression. (A) *Igf2* expression is high in neonatal hepatocytes (postnatal days 1-7, P1-7) but becomes undetectable in matured hepatocytes (P21 and P56), closely correlating with the cell cycle signature and YAP/TAZ activity. The transcriptome of postnatal hepatocytes was analyzed using scRNA-seq data from developing livers (GSE171993). (B and C) *Igf2* expression is tightly associated with canonical YAP/TAZ target genes in mouse livers with Hippo pathway perturbations at different ages. Gene Set Enrichment Analysis (B) and heatmap (C) are shown. The RNA-seq data for *YAPTg* livers were analyzed using data from GSE178227. (D) Volcano plot displaying differentially expressed genes in 16-month-old *Sav1^−/−^;Wwc*^−/−^ livers compared to WT livers, with genes of interest marked. (E) Schematic diagram illustrating the experimental procedure. Primary hepatocytes were isolated from *Rosa26-tdTomato;Sav1^F/F^;Wwc^F/F^* mice, treated with Ad-*Cre* and siRNA after cell attachment, and analyzed three days post-isolation. In *Rosa26-tdTomato* mice, tdTomato expression labels Cre-expressing cells. (F and G) *IGF2* expression is elevated in primary hepatocytes lacking *Sav1* and *Wwc1/2*, but decreases with simultaneous knockdown of *Yap*/*Taz*. Quantitative RT-PCR results (F) and immunostaining images (G) of IGF2 (green), tdTomato (red), and nuclei (blue) are shown. Scale bar in G: 10 μm. (H-J) *LATS1/2* knockout in HEPG2 cells induces *IGF2* expression, while *LATS1/2* overexpression inhibits *IGF2* expression. Immunoblotting (H), immunostaining images (I) of IGF2 (red) and nuclei (blue), and quantitative RT-PCR results (J) are shown. Scale bar in I: 10 μm. (K) Knockdown of *Tead1-4* in primary mouse hepatocytes with *Sav1;Wwc1/2* deletion impairs *Igf2* and *Cyr61* expression. Data in F, J, and K are presented as mean ± SEM from at least three independent biological replicates, and *P* values were assessed by unpaired Student’s t-tests. **p < 0.01, ***p < 0.001, ****p < 0.0001; ns: not significant.

**Figure S2.**
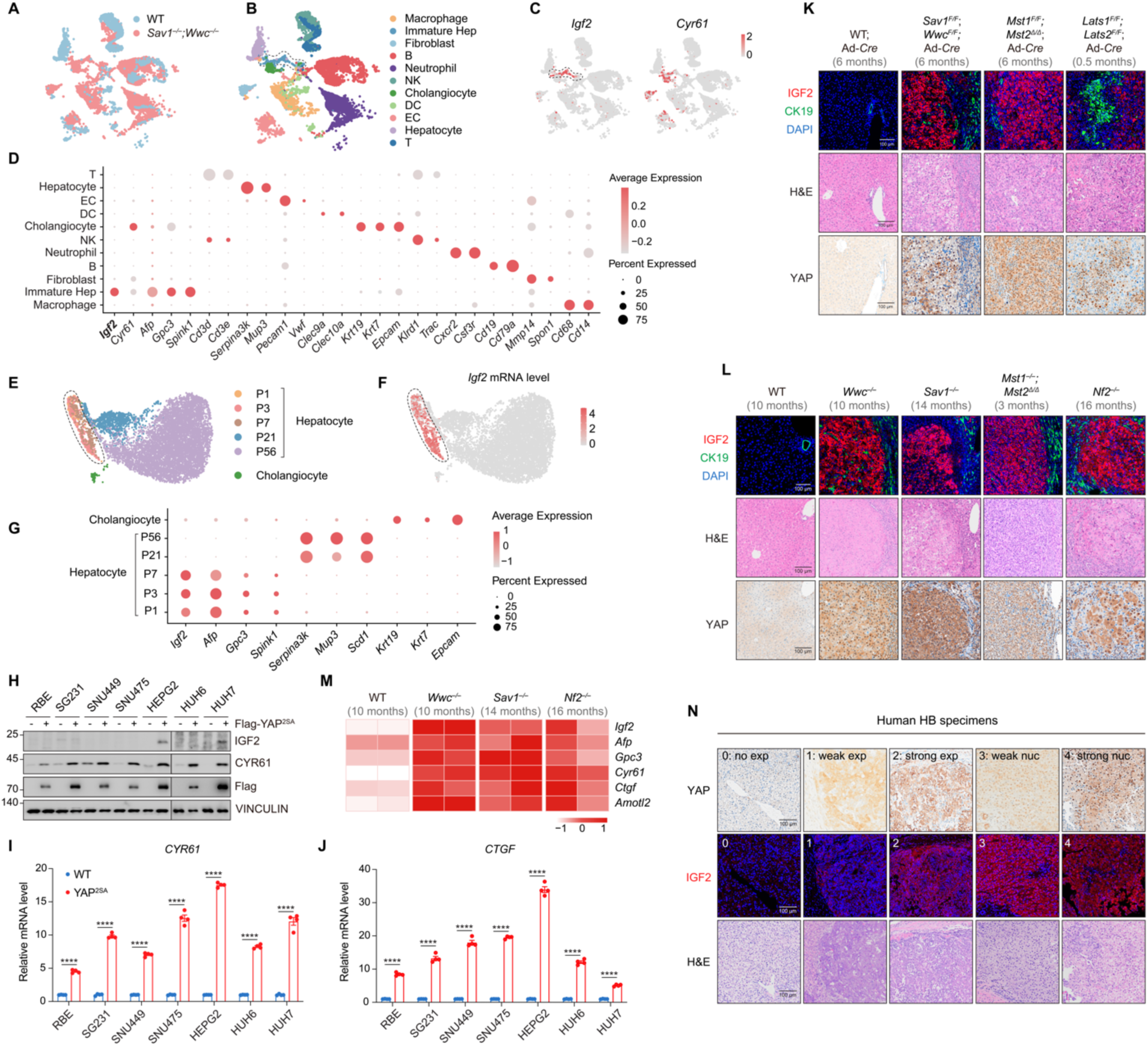
IGF2 is regulated by the Hippo pathway in immature hepatocytes and hepatoblastomas. (A and B) UMAP representation of integrated liver cells, generated by combining scRNA-seq data from 3-week-old WT (GSE171993) and *Sav1^−/−^;Wwc*^−/−^ livers. Cells are colored by samples (A) or cell types (B). Immature Hep: immature hepatocytes; B: B cells; NK: natural killer cells; DC: dendritic cells; EC: endothelial cells; T: T cells. (C) Expression profiles of *Igf2* and *Cyr61* in liver cells. (D) Expression of selected markers for different cell types. Dot size and color indicate the proportion of cells expressing marker genes in each cluster and the average gene expression level, respectively. (E) UMAP representation of integrated scRNA-seq data from developing mouse livers (GSE171993). (F) Expression profile of *Igf2* in developing mouse livers. (G) Expression of *Igf2*, immature hepatocyte markers (*Afp*, *Gpc3*, *Spink1*), matured hepatocyte markers (*Serpina3k*, *Mup3*, *Scd1*), and cholangiocyte markers (*Krt19*, *Krt7*, *Epcam*). Dot size and color indicate the proportion of cells expressing marker genes and the average gene expression level. (H) YAP^2SA^ effectively induces *IGF2* expression in HEPG2 and HUH7 cells but not in cells where *IGF2* is silenced. (I and J) YAP^2SA^ effectively induces *CYR61* and *CTGF* expression in diverse cancer cells. Data are presented as mean ± SEM from at least three independent biological replicates. *P* values were assessed by unpaired Student’s t-tests. ****p < 0.0001. (K and L) IGF2 is highly expressed in YAP/TAZ-activated HCC but not ICC or bile duct hamartomas surrounding HCC. H&E staining and immunostaining of IGF2 (red), CK19 (green), nuclei (blue), and YAP (IHC) are shown. Scale bar: 100 μm. (M) HCC driven by dysregulated Hippo pathway exhibits immature hepatocyte features, as indicated by the expression of *Igf2*, immature hepatocyte markers (*Afp*, *Gpc3*, *Spink1*), and canonical YAP/TAZ target genes (*Ctgf*, *Amotl2*). (N) H&E staining and immunostaining of IGF2 (red) and YAP (IHC) in human HB specimens. Each column represents data from the same patient. YAP levels were graded based on both expression and nuclear localization: 0, no expression; 1, weak expression; 2, strong expression; 3, weak nuclear localization; 4, strong nuclear localization. Scale bar: 100 μm.

**Figure S3.**
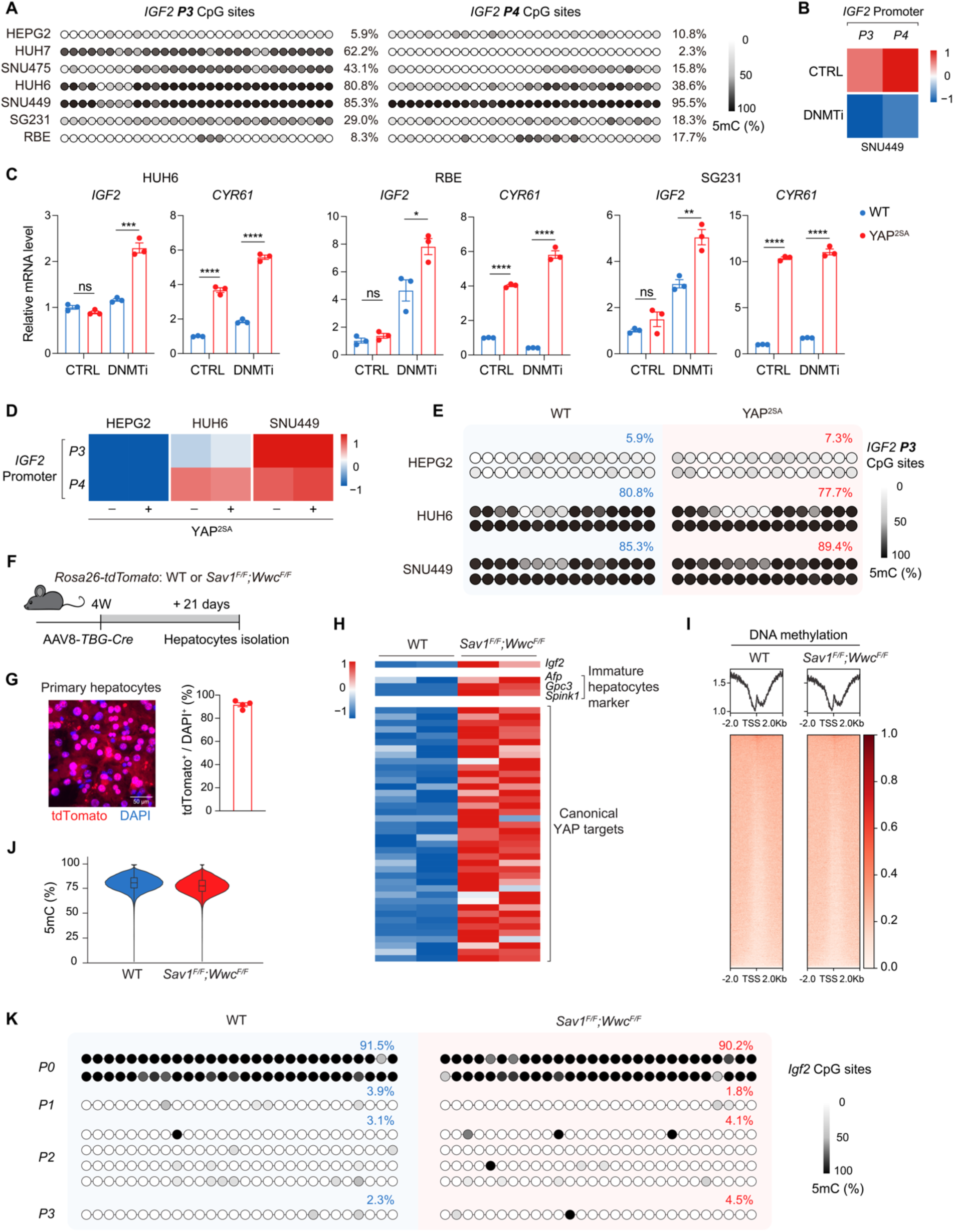
YAP/TAZ-induced *IGF2* expression is impeded by promoter DNA methylation. (A) DNA 5mC levels at the *IGF2 P3* (left) and *P4* (right) fetal promoters. Each dot represents a CpG site within the *IGF2 P3/4* region, and the color reflects the 5mC level at that site. (B) The DNA methyltransferase inhibitor (DNMTi) decitabine treatment reduces 5mC levels at *IGF2 P3/P4* promoters in SNU449 cells. (C) *IGF2* expression is reactivated and further enhanced by YAP^2SA^ when treated with decitabine in HUH6, RBE, and SG231 cells. Data are presented as mean ± SEM from at least three independent biological replicates. *P* values were assessed by unpaired Student’s t-tests. *p < 0.05, **p < 0.01, ***p < 0.001, ****p < 0.0001; ns: not significant. (D) DNA methylation levels at the *IGF2 P3/P4* promoters remain largely unchanged upon ectopic expression of YAP^2SA^ in HEPG2, HUH6, and SNU449 cells. (E) YAP^2SA^ expression has no obvious effect on DNA methylation at the *IGF2 P3* promoter. Each dot represents a CpG site, and the color reflects the 5mC level at this site. (F) Schematic diagram illustrating the experimental procedure. *Rosa26-tdTomato;Sav1^F/F^;Wwc^F/F^*mice and *Rosa26-tdTomato*;WT mice received tail vein injection of AAV8-*TBG-Cre* to induce hepatocyte-specific gene knockout at 4 weeks old. Primary hepatocytes were then isolated 21 days post-injection. (G) Analysis of tdTomato expression indicates that the purity of the isolated primary hepatocytes and the efficiency of Cre recombinase were approximately 90%. Immunostaining images (left) of tdTomato (red) and nuclei (blue) and quantifications (right) are shown. Scale bar: 50 μm. (H) Transcriptome analysis of WT and *Sav1^F/F^;Wwc^F/F^* hepatocytes. (I and J) Global 5mC levels exhibit minimal differences between WT and *Sav1^F/F^;Wwc^F/F^* hepatocytes. Violin plot (I) and heatmap (J) centered at transcription start sites (TSS) are shown. (K) DNA methylation levels at the *Igf2 P0-P3* promoters in WT and *Sav1^F/F^;Wwc^F/F^*hepatocytes. Each dot represents a CpG site within the promoter region, and the color reflects the 5mC level at this site.

**Figure S4.**
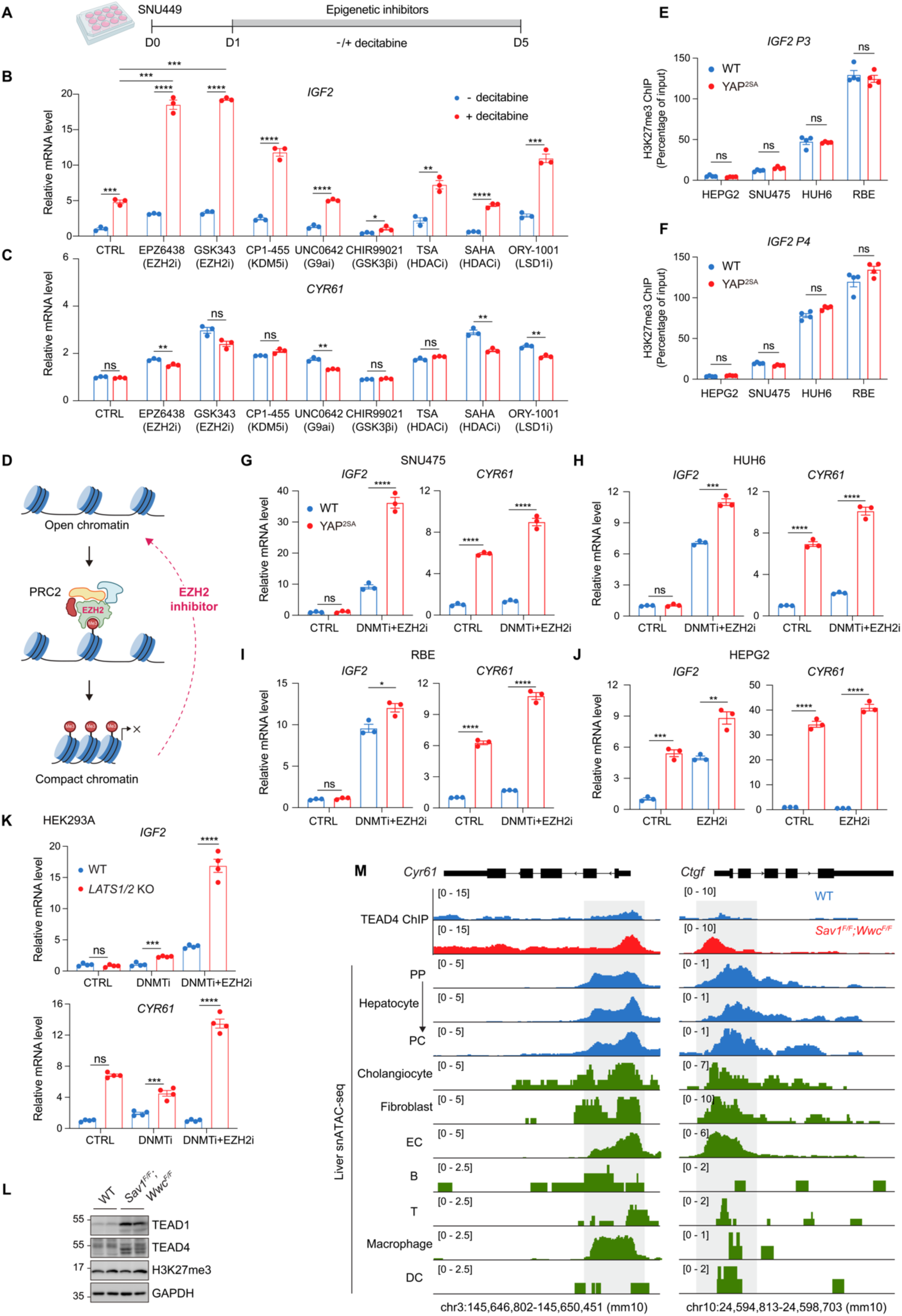
YAP/TAZ-induced *IGF2* expression is prevented by promoter histone methylation. (A) Schematic diagram illustrating the experimental procedure. SNU449 cells were treated with various epigenetic inhibitors (5 μM) for 96 hours in the presence or absence of the DNA methyltransferase inhibitor (DNMTi) decitabine (50 μM). (B and C) Quantitative RT-PCR results of *IGF2* (B) and *CYR61* (C) in SNU449 cells when treated with various epigenetic inhibitors. Blue bars represent the absence of decitabine, and red bars represent the presence of decitabine. (D) Schematic diagram illustrating how H3K27me3 functions as a gene silencer through chromatin compaction. H3K27me3 is catalyzed by EZH2, the enzymatic component of Polycomb Repressive Complex 2 (PRC2). EZH2 inhibitors (EZH2i) can reduce H3K27me3 levels, leading to chromatin open and gene activation. (E and F) YAP does not actively remove H3K27me3 modifications near *IGF2* promoters in liver cancer cells. ChIP signals were normalized to input DNA. (G-K) Treatment with the EPZ6438, in combination with decitabine, enhances YAP-induced *IGF2* expression in SNU475, HUH6, RBE, HEPG2, and HEK293A cells. (L) Immunoblotting indicates that deletion of *Sav1;Wwc1/2* in hepatocytes increases TEAD1/4 expression, while H3K27me3 remains unchanged. (M) Genomic tracks displaying TEAD4 ChIP-seq and mouse liver snATAC-seq (González-Blas et al. 2024) reads at the *Cyr61* (left) and *Ctgf* (right) loci. TEAD-binding sites are marked in gray, which are accessible in hepatocytes, cholangiocytes, fibroblasts, ECs, and several other mesenchymal cells. PP: periportal; PC: pericentral; EC: endothelial cell; DC: dendritic cell. Data in B, C, and E-K are presented as mean ± SEM from at least three independent biological replicates. *P* values were assessed by unpaired Student’s t-tests. *p < 0.05, **p < 0.01, ***p < 0.001, ****p < 0.0001; ns: not significant.

**Figure S5.**
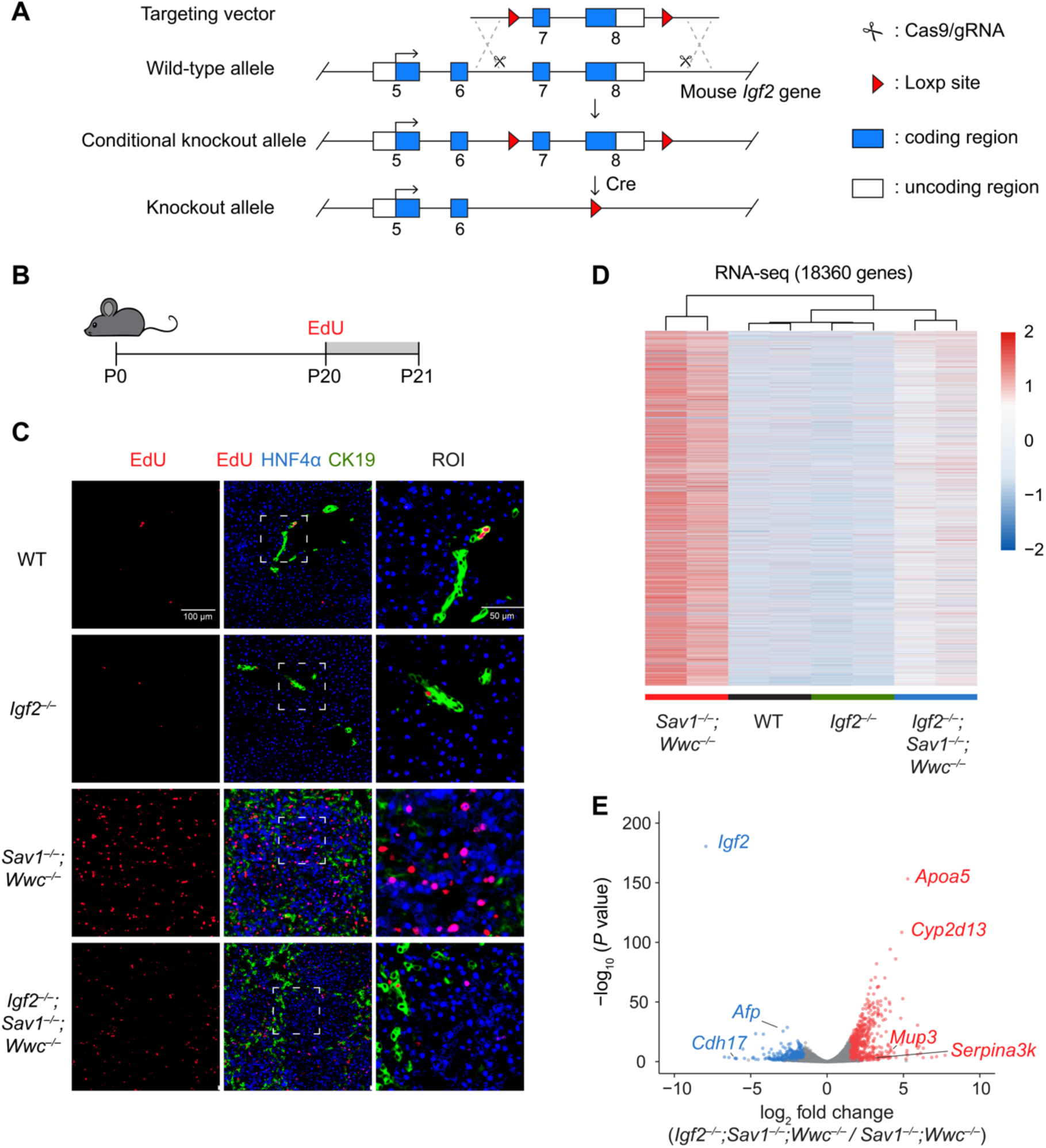
IGF2 deficiency impairs YAP/TAZ-induced liver enlargement and HB development. (A) Strategy for generating *Igf2* conditional knockout mice. Exon 7 and exon 8 of *Igf2* were targeted, which contain the functional coding sequence shared by all *Igf2* transcripts. The targeting vector was electroporated into JM8A3 embryonic stem cells. Positive clones were confirmed by PCR and injected into C57BL/6J mouse blastocysts to generate chimeric mice. Exon 7 and exon 8 were removed upon Cre-mediated recombination. (B) Schematic diagram illustrating the procedure of *in-situ* EdU labeling. EdU (50 mg/kg) was injected intraperitoneally 24 hours before sacrifice. (C) Staining images of EdU (red), HNF4α (blue), and CK19 (green). Scale bar: 100 μm. Region of interest (ROI) scale bar: 50 μm. (D) Clustering analysis of transcriptomes form 3-week-old livers. (E) Volcano plot displaying differentially expressed genes in 3-week-old *Igf2^−/−^;Sav1^−/−^;Wwc*^−/−^ livers compared to *Sav1^−/−^;Wwc*^−/−^ livers, with genes of interest marked.

**Figure S6.**
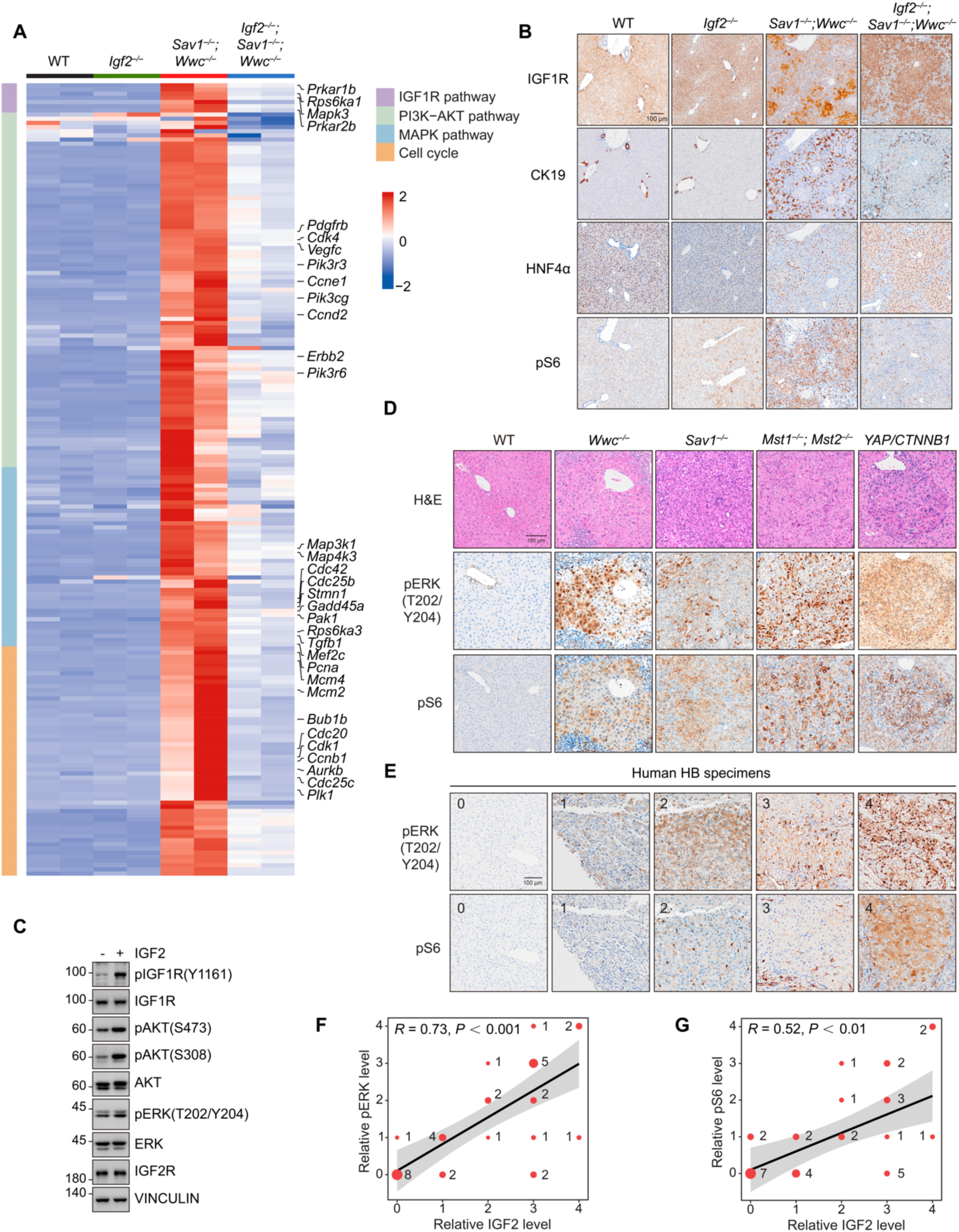
Activation of YAP/TAZ-IGF2-IGF1R signaling in liver growth and tumorigenesis. (A) Heatmap showing the expression profiles of core enriched genes in the IGF1R pathway, PI3K-AKT pathway, MAPK pathway, and cell cycle in 3-week-old livers. Genes associated with cell proliferation are indicated on the right side. (B) Immunostaining of IGF1R, CK19, HNF4α, and pS6 in liver sections from 3-week-old WT, *Igf2^−/−^*, *Sav1^−/−^;Wwc*^−/−^, and *Igf2^−/−^;Sav1^−/−^;Wwc*^−/−^ mice. Scale bar: 100 μm. (C) Activation of IGF1R signaling in HEPG2 cells. Purified human IGF2 protein (500 ng/mL) was added to the culture medium of HEPG2 cells for 1 hour. After incubation, cells were lysed and subjected to immunoblotting. (D) Immunostaining of pERK (T202/Y204) and pS6 in IGF2-expressing liver tumors with dysregulated Hippo signaling. Scale bar: 100 μm. (E) Immunostaining of pERK (T202/Y204) and pS6 in human HB specimens. Immunostaining images of IGF2 and YAP are shown in Figure S2N. Each column represents data from the same patient. Scale bar: 100 μm. (F and G) Quantifications of pERK (T202/Y204) (F) and pS6 (G) in human HB specimens. Dot size indicates the number of specimens. Correlation coefficients were analyzed using Spearman’s correlation.

**Figure S7.**
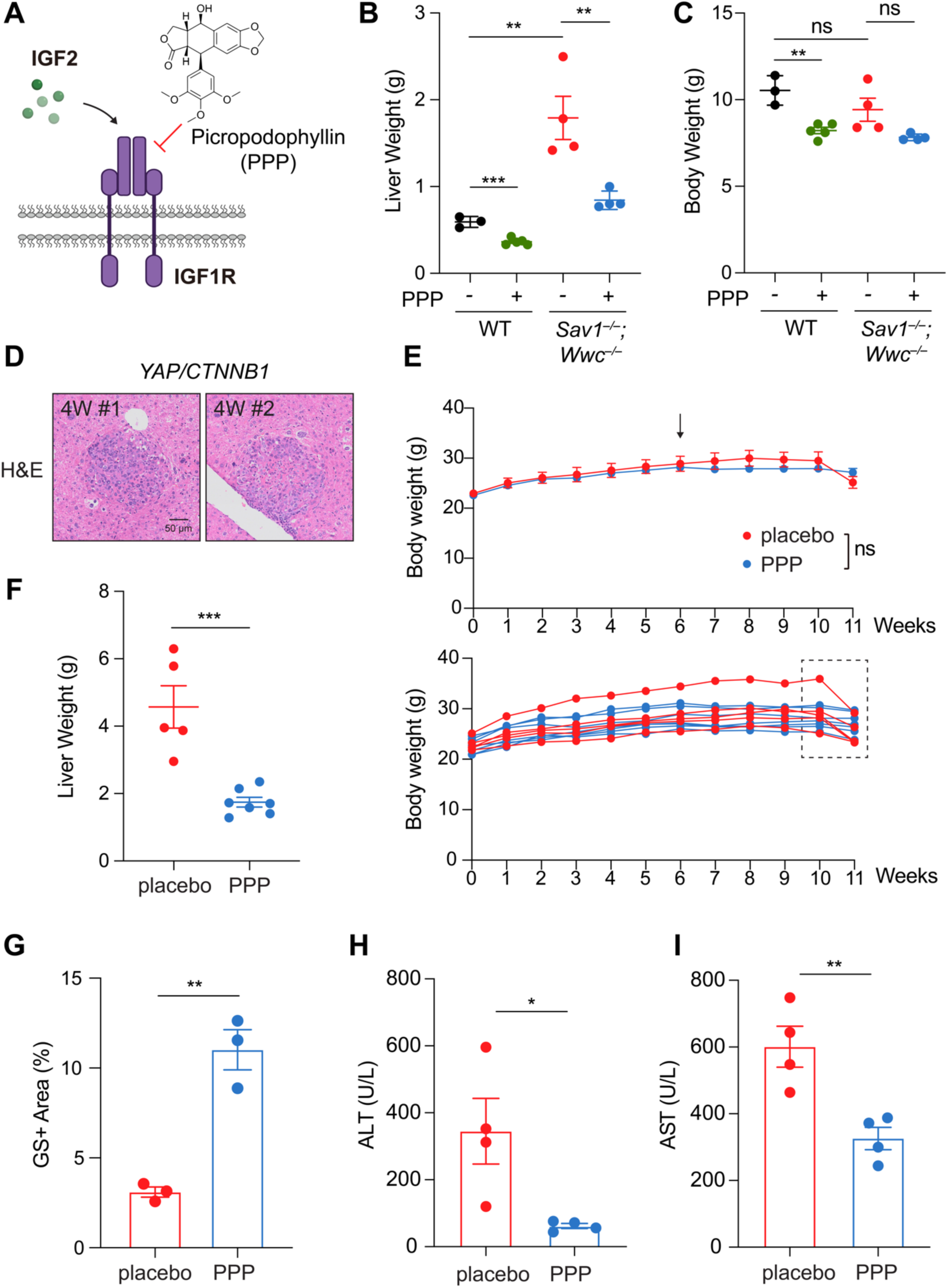
IGF1R inhibition prevents YAP/TAZ-induced liver enlargement and hepatoblastoma. (A) Schematic diagram illustrating the structure and target of Picropodophyllin (PPP). (B and C) Liver weight (B) and body weight (C) of 3-week-old WT and *Sav1^−/−^;Wwc*^−/−^ mice, treated with either PPP or placebo. (D) HB tumor lesions were observed by 4 weeks post hydrodynamic tail vein injections (HDI) of *YAP/CTNNB1*. H&E staining images are shown, with numbers representing different mice. (E) Body weight of mice treated with placebo (red) or PPP (blue). Data are presented as mean ± SEM (upper panel) or individually for each mouse (lower panel). Rapid weight loss was observed in placebo-treated mice around 11 weeks post-HDI (dashed box). Statistical significance was determined using one-way analysis of variance (ANOVA). (F) Liver weight of mice treated with placebo (red) or PPP (blue). (G) Quantification results of GS staining in Figure 7I. (H and I) Alanine aminotransferase (ALT) and aspartate aminotransferase (AST) levels of mice treated with placebo or PPP. Data in B, C, and E-I are presented as mean ± SEM, and each point represents an individual mouse. *P* values were assessed by unpaired Student’s t-tests. *p < 0.05, **p < 0.01, ***p < 0.001; ns: not significant.

**Figure S8.**
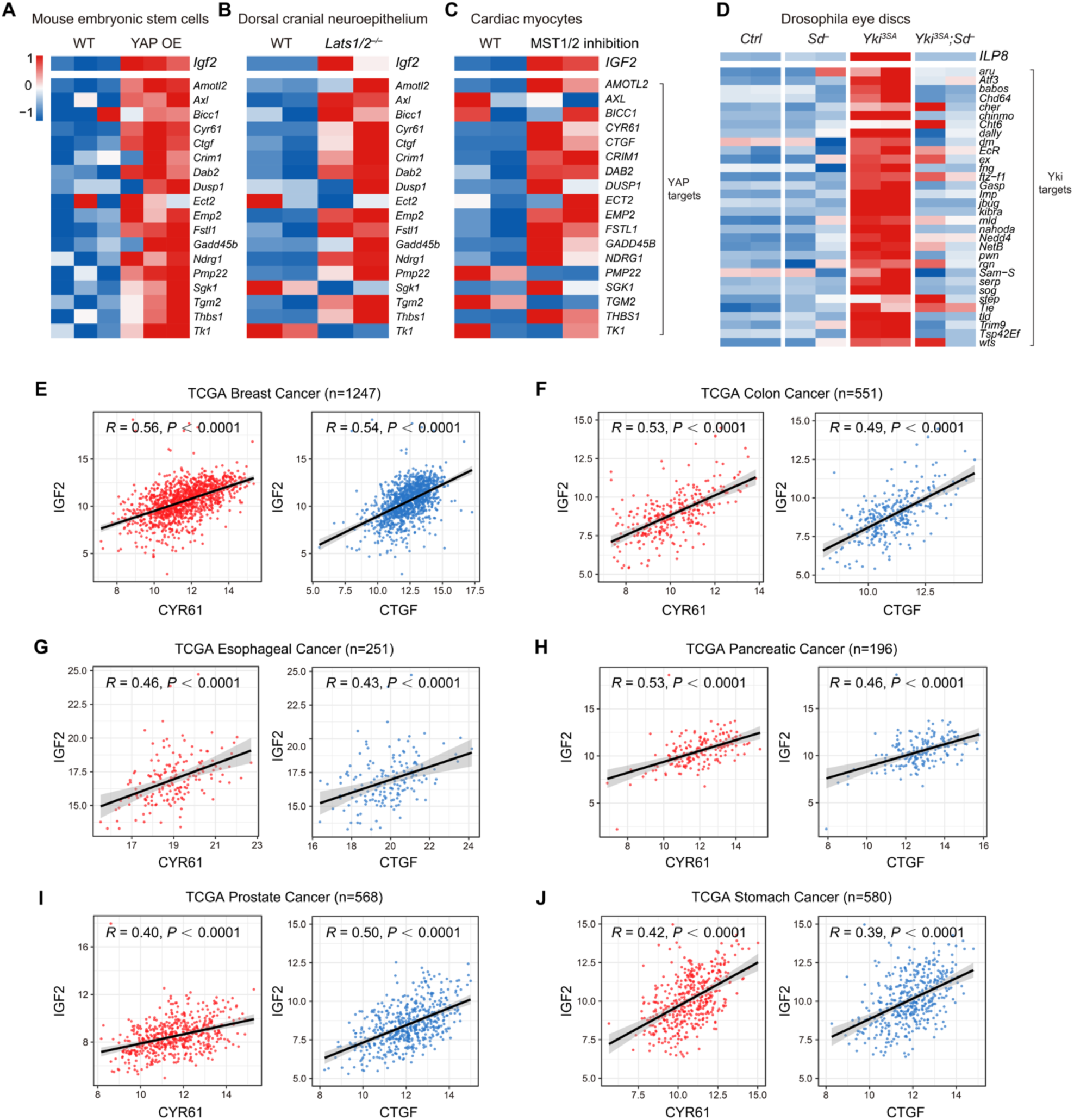
Conservation of Hippo-IGF2 axis across diverse organs, species, and cancers. (A-C) Under diverse conditions that lead to YAP activation, *IGF2* expression is simultaneously upregulated along with other canonical YAP/TAZ target genes in mouse embryonic stem cells (GSE157706) (A), dorsal cranial neuroepithelium (GSE182721) (B), and cardiac myocytes (GSE176142) (C). (D) The expression of *insulin-like peptide 8* (*ILP8*) is regulated by Yorkie (the ortholog of YAP/TAZ) in a Scalloped (Sd, the ortholog of TEAD)-dependent manner in *Drosophila* eyes (GSE211458). (E-J) *IGF2* expression is highly associated with canonical YAP/TAZ target genes in breast cancer (E), colon cancer (F), esophageal cancer (G), pancreatic cancer (H), prostate cancer (I), and stomach cancer (J). Correlation coefficients were analyzed using Spearman’s correlation.

